# Comparison of two individual-based model simulators for HIV epidemiology in a population with HSV-2 using as case study Yaoundé-Cameroon, 1980-2005

**DOI:** 10.1101/637389

**Authors:** Diana M Hendrickx, João Dinis Sousa, Pieter J.K. Libin, Wim Delva, Jori Liesenborgs, Niel Hens, Viktor Müller, Anne-Mieke Vandamme

## Abstract

Model comparisons have been widely used to guide intervention strategies to control infectious diseases. Agreement between different models is crucial for providing robust evidence for policy-makers because differences in model properties can influence their predictions. In this study, we compared models implemented by two individual-based model simulators for HIV epidemiology in a population with Herpes simplex virus type 2 (HSV-2). For each model simulator, we constructed four models, starting from a simplified basic model and stepwise including more model complexity. For the resulting eight models, the predictions of the impact of behavioural interventions on the HIV epidemic in Yaoundé (Cameroon) were compared. The results show that differences in model assumptions and model complexity can influence the size of the predicted impact of the intervention, as well as the predicted qualitative behaviour of the HIV epidemic after the intervention. Moreover, two models that agree in their predictions of the HIV epidemic in the absence of intervention can have different outputs when predicting the impact of interventions. Without additional data, it is impossible to determine which of these two models is the most reliable. These findings highlight the importance of making more data available for the calibration and validation of epidemiological models.

## Introduction

Mathematical modelling has been widely applied to better understand the transmission, treatment and prevention of infectious diseases. The role of mathematical models in understanding the dynamics of the human immunodeficiency virus (HIV) epidemic has been recently reviewed by Geffen & Welte^1^. They present examples of HIV models that were used for estimating the size of the epidemic in specific subpopulations such as the Black population in South Africa and for estimating the impact of interventions such as the use of antiretrovirals and condoms to reduce HIV and AIDS.

When considering mathematical models, two commonly used types of implementation can be distinguished: compartmental and individual-based models (IBMs). While compartmental models simulate population counts, IBMs (also called agent-based models or micro-simulation models) keep track of the history of each individual in the population separately.

Recent applications of compartmental HIV models include testing the effect of different assumptions for HIV dynamics on the predicted impact of antiretroviral therapy (ART) in men-having-sex-with-men (MSM)^2^, and the study of the influence of concurrent partnerships on HIV dynamics^3^. IBMs have been recently applied to assess the influence of pre-exposure prophylaxis (PrEP) in MSM^4^, to evaluate the long-term effect of early ART initiation^5^ and to understand the factors underlying the emergence of HIV in humans^6^.

In contrast to compartmental models, IBMs allow a high degree of individual heterogeneity^7^ (e.g. regarding (sexual) risk behaviour, age demography and individual response to treatment). This is an advantage when heterogeneity matters for the particular question/process that is studied. Because individual heterogeneity is inherent in transmission, prevention and treatment of HIV and other sexually transmitted infections (STI), IBMs are particularly suitable to estimate the most beneficial intervention for specific individuals. Furthermore, IBMs allow for explicit modelling of the sexual relationships that jointly form the sexual network over which STI/HIV infections are transmitted^8^.

Model comparisons are crucial for providing robust evidence for decision-making in public health and policy purposes^9^. To assess uncertainty, it is necessary to study how differences in model properties influence their prediction. Eaton et al. compared ten mathematical models aimed at studying HIV prevalence, incidence and ART^10^. However, in contrast to the models in our study, eight of these models were compartmental, and the majority of these models did not include the co-factor effects of other STIs. The remaining two models were discrete-time IBMs.

This study presents and compares eight models generated with two IBM frameworks for simulating HIV transmission dynamics with STI co-factor effects. The purpose of the comparison was to assess how differences in properties between the eight models influence the prediction of the impact of behavioural interventions on the HIV epidemic. For this purpose, we chose the HIV epidemic in Yaoundé (Cameroon) during the period 1980–2005, and calibrated all models to HIV prevalence time series from Yaoundé.

## Methods

### Individual-based models

In this study, individual-based models developed with the Simpact Cyan 1.0 and StepSyn 1.0 modelling frameworks, were compared. A short description of each modelling framework is given below. More details about differences between the two modelling frameworks and the generation of the sexual networks are provided in the Supplementary Material, Section 1. For each modelling framework, four models were generated, starting from a simplified basic model and incorporating additional complexity.

#### Simpact Cyan 1.0 modelling framework

Simpact Cyan 1.0 (http://www.simpact.org/)^11^ is a freely available framework for developing IBMs to simulate HIV transmission, progression and treatment. Models developed with Simpact Cyan 1.0 are event-driven, which means that the models are not updated at fixed time intervals, but at every time an event happens, making Simpact Cyan 1.0 a continuous-time simulation modelling framework. Events occur as a result of stochastic event processes, described by hazard functions. The timing of an event is determined using the modified next reaction method (mNRM)^12^.

To initialize a model, a number of individuals are generated. The age of each person is drawn from an age distribution based on a population pyramid, and when the age is larger or equal to the sexual debut age, a person is marked as sexually active.

In Simpact Cyan 1.0, the user can specify which events are possible in the simulation, depending on the research question he or she wants to answer. The possible events are heterosexual and homosexual relationship formation and dissolution, conception and birth, HIV transmission, AIDS and non-AIDS mortality, HIV diagnosis and treatment, HIV treatment dropout, and STI transmission events (in this study HSV-2). Furthermore, events describing the natural history of HIV are implemented in Simpact Cyan 1.0. For each event, the form and the parameters of the hazard function can be modified flexibly. The HIV transmission hazard can be described in terms of an individual’s viral load. In the basic model in this study, this option, as well as age-dependency, is turned off. Simpact Cyan 1.0 is implemented in C++ with R^13^ and Python^14^ interfaces.

#### StepSyn 1.0 modelling framework

StepSyn 1.0 is an IBM modelling framework that simulates epidemics of STIs, including HIV. It focuses on the epidemiological synergy and interactions between HIV and other STIs. Time is divided into fixed one-week intervals. Formation and dissolution of sexual relationships and STI and HIV transmission are modelled as stochastic processes. For each STI, life history is explicitly modelled, including several stages, symptomatic recurrences and genital ulcers. The effects of these symptoms on the HIV transmission probability are explicitly modelled. The parameters for the timing and duration of the STI stages and symptoms and the co-factor effects of genital ulcers on HIV transmission are based on a literature review. In the present study, StepSyn 1.0 is parameterized to track only HIV and Herpes simplex virus type 2 (HSV-2). In the basic model the latter virus is modelled without explicit stages or symptomatic recurrences, and with the probability of transmission and co-factor effect on HIV both constant and not dependent on recurrences. In this study, StepSyn is ran in both this basic model, and in the full model in which HSV-2 stages and recurrences are modelled. Heterosexual relationships can include marital and short-term links and contacts between commercial sex workers (CSW) and clients. In the present study, CSWs are not included. Individuals vary in their tendency to form short-term relationships so that their number of non-marital partners within a year would follow a power-law distribution. The parameters of the latter were derived from behavioural data gathered in the 4 Cities Study^15,16^. StepSyn is implemented in R^13^. StepSyn is not yet freely available but will become so upon publication of a manuscript that describes the modelling framework into more detail (manuscript in preparation).

#### Models used in this study

The following models were used in this study:

1. Simpact Cyan 1.0 basic model (Si_Ba). This simplified model assumes no population inflow and no outflow by other causes than AIDS. Furthermore, HIV transmission doesn’t depend on an individual’s viral load. HSV-2 is implemented as a generic STI co-factor, without modelling the natural history. This means that only the impact of HSV-2 on HIV acquisition and transmission, and vice versa, are implemented.
2. Simpact Cyan 1.0 model with population inflow and outflow (Si_IO). Birth is implemented as an event related to relationship formation, followed by a conception event, using the default settings of Simpact Cyan 1.0. Non-AIDS mortality is age-dependent and is derived from the 1980 population pyramid of Yaoundé ^17^. Other settings are the same as in the Si_Ba model.
3. Simpact Cyan 1.0 model implementing viral load (Si_VL). HIV transmission depends on an individual’s viral load. Other settings are the same as in the Si_Ba model.
4. Simpact Cyan 1.0 model implementing population inflow, outflow and viral load (Si_IO_VL). Birth and non-AIDS mortality are implemented in the same way as for the Si_IO model. HIV transmission depends on an individual’s viral load.
5. StepSyn 1.0 basic model (St_Ba). This simplified model assumes no population inflow (birth, immigration) and no outflow by other causes than AIDS. Furthermore, HSV-2 is modelled without explicit stages or recurrences, and with the probability of transmission and co-factor effect on HIV both constant and not dependent on recurrences.
6. StepSyn 1.0 model with population inflow and outflow (St_IO). This model implements birth, non-AIDS mortality, immigration and emigration. All of these variables are expressed as a rate. Other settings are the same as in the St_Ba model.
7. StepSyn 1.0 model with the full set of HSV-2 co-factor assumptions (St_RG). HSV-2 life history is explicitly modelled, including several stages, recurrences and genital ulcers. The effects of these symptoms on the HIV transmission probability are explicitly modelled. Other settings are the same as in the St_Ba model.
8. StepSyn 1.0 model with population inflow, outflow and the full set of HSV-2 co-factor assumptions (St_IO_RG). This model implements birth, immigration, non-AIDS mortality and emigration in the same way as the Si_IO model. For HSV-2, the same settings as in the St_RG model are used.

### Calibration of the models

For the eight models, HIV transmission parameters were fitted to HIV prevalence data of Yaoundé from 1989 to 1998. A detailed description of the data is provided in the Supplementary Material, section 2. The parameter estimation procedure is described in section 3 of the Supplementary Material.

Furthermore, we adapted the parameters for STI (HSV-2) transmission. In major epidemiological reviews of HSV-2^18–20^ the only data about Yaoundé that is mentioned is the data collected by Buvé et al.^21^ in the 4 Cities Study, in which the HSV-2 seroprevalence was 50% in females. Thus, we do not have information about the temporal trends of HSV-2 in Yaoundé. However, since the prevalence in Africa seems to have slightly declined between 2003 and 2012^18^ we assumed that the measured prevalence in Yaoundé, in 1997 ^21^, reflects an epidemic not far from its peak. Accordingly, we adapted the HSV-2 transmission parameters of the eight models so that the seroprevalence of this virus first increases and afterwards stabilizes at approximately 50% for females, corresponding to the HSV-2 prevalence in 1997 described by Buvé et al.^21^.

The estimated parameters and other key parameters for each of the eight models are described in the Supplementary Material, section 4.

All models could be calibrated so that they fit the HIV prevalence data between 1989 and 1998, and the HSV-2 prevalence first increases and afterwards stabilizes at approximately 50% for females (see Supplementary Material, section 5).

### Validation of the models against data not used for fitting

The eight models were validated against the HIV prevalence in 2004, which was not used for model calibration and has been reported to be 6.0% in males and 10.7% in females^22^. For each of the eight models, we checked how well these literature values could be predicted.

### Prediction of the impact of behavioural interventions

To investigate the impact of behavioural interventions on the HIV epidemic, the distribution of the number of partners was changed in all of the eight models to study the impact of behavioural change with respect to promiscuity. Only a single parameter was changed, we refer to this parameter as “behavioural change parameter”. More details on the behavioural change parameter for Simpact Cyan 1.0 and StepSyn 1.0 are provided in section 6 of the Supplementary Material.

Changing the behavioural change parameter in Simpact Cyan 1.0 and StepSyn 1.0, as described in section 6 of the Supplementary Material has a similar effect on the distribution of the number of partners in both modelling frameworks:

- the mean number of partners is reduced by 6%;
- the 95% percentile is reduced with 1;
- the median and 75% percentile are unchanged.

To investigate what would have happened to the epidemic if promiscuity would have been lower from day 1 onwards, we implemented the change of the behavioural change parameter described above in 1980.

Second, the intervention was applied in 1990 to study its impact when it would have been implemented at some point in time during the epidemic.

When studying the impact of interventions, only the parameter(s) related to the interventions (in this study only the behavioural change parameter) are changed. This means that we assume that apart from the intervention, no other influences are present. Therefore, the impact of behavioural interventions was also explored during the study period 1980-2005.

## Results

### Validation of the models against data not used for fitting

The median HIV prevalence from 1980 to 2005 (in %), together with the literature value for the HIV prevalence in 2004^22^ is shown in Fig. 1. All models simulating no population inflow and outflow, except the StepSyn 1.0 model with STI life history explicitly modelled (St_RG), largely overestimate the HIV prevalence in 2004 for both females and males. In contrast, the St-RG model underestimates the HIV prevalence in 2004 and the HIV prevalence stabilizes after 2000 and 2004 for females and males respectively. Implementing inflow and outflow considerably improves the prediction of the HIV prevalence for both females and males in 2004. For females, the Simpact 1.0 model including inflow, outflow and a VL-dependent HIV transmission hazard (Si_IO_VL) results in the prediction closest to the literature value. For males, the StepSyn 1.0 model with inflow, outflow and STI life history explicitly modeled (St_IO_RG) results in the best prediction of the value in the literature. In general, the StepSyn 1.0 models showed less variation between simulations in predicted HIV prevalence in 2004 than the Simpact 1.0 models (Table 1).

**Figure 1.**
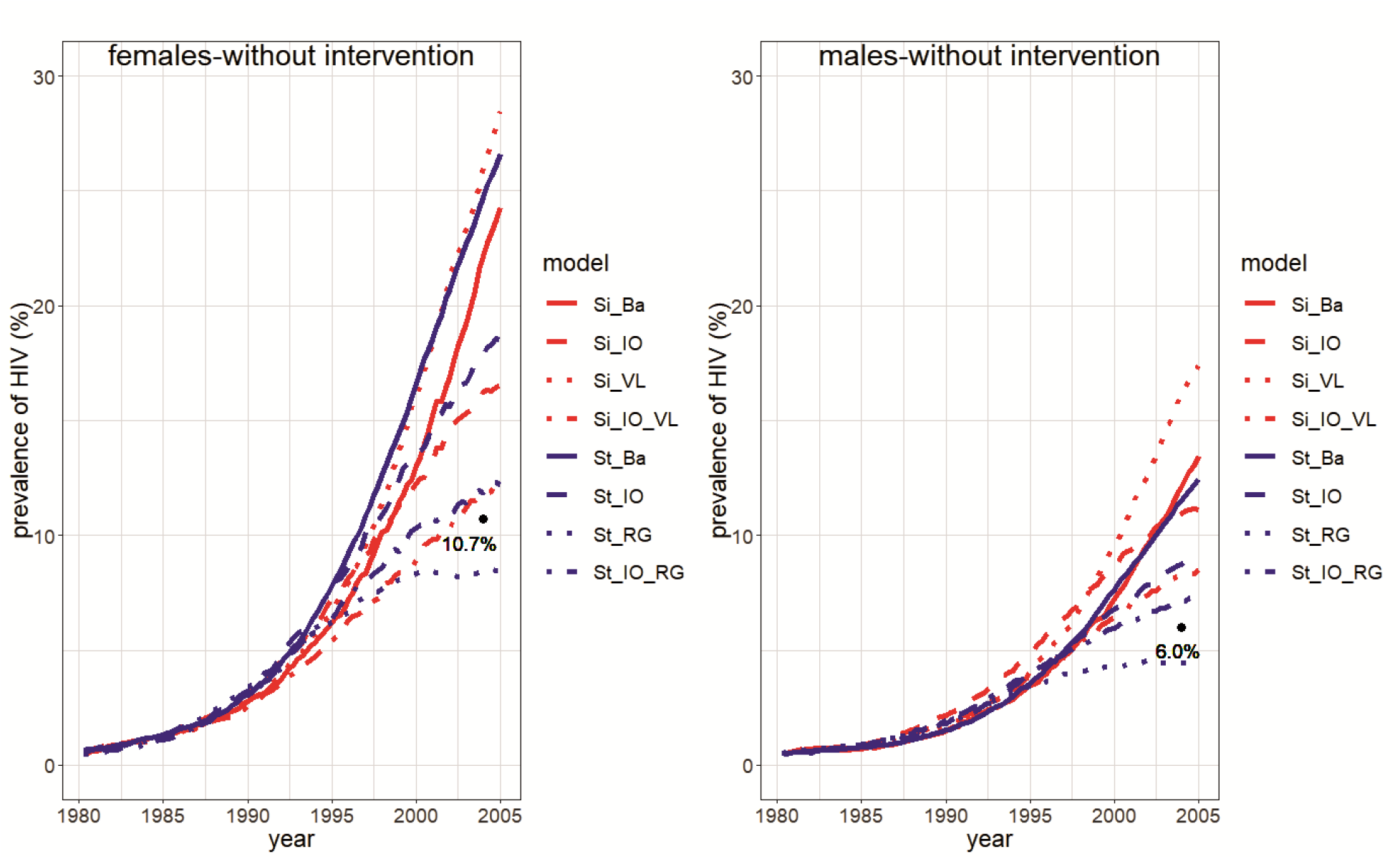
Prevalence curves for HIV from 1980 to 2005 in case no behavioural intervention is implemented. Median HIV prevalence (in %) of 100 simulations. Left: females; right: males. Models: Si_Ba: Simpact 1.0 basic model; Si_IO: Simpact 1.0 model with inflow and outflow; Si_VL: Simpact 1.0 model with VL-dependent HIV transmission hazard; Si_IO_VL: Simpact 1.0 model with inflow, outflow, VL-dependent HIV transmission hazard; St_Ba: StepSyn 1.0 basic model; St_IO: Stepsyn 1.0 model with inflow and outflow; St_RG: StepSyn 1.0 model with STI life history explicitly modeled; St_IO_RG: StepSyn 1.0 model with inflow, outflow and STI life history explicitly modeled. The black dot represents the literature value from ^22^.

**Table 1.** Results for the predicted HIV prevalence in 2004 for females and males in case of no behavioural intervention, a behavioural intervention implemented in 1990, and a lower promiscuity from 1980 onwards. Median HIV prevalence (in %) and range ([minimum,maximum]) of 100 simulations. Models: Si_Ba: Simpact 1.0 basic model; Si_IO: Simpact 1.0 model with inflow and outflow; Si_VL: Simpact 1.0 model with VL-dependent HIV transmission hazard; Si_IO_VL: Simpact 1.0 model with inflow, outflow, VL-dependent HIV transmission hazard; St_Ba: StepSyn 1.0 basic model; St_IO: Stepsyn 1.0 model with inflow and outflow; St_RG: StepSyn 1.0 model with STI life history explicitly modeled; St_IO_RG: StepSyn 1.0 model with inflow, outflow and STI life history explicitly modeled.

| Model | No intervention |  | Intervention in 1990 |  | Lower promiscuity from 1980 onwards |  |
| --- | --- | --- | --- | --- | --- | --- |
|  | Females | Males | Females | Males | Females | Males |
| Si_Ba | 22.1 (6.3-34.5) | 12.1 (3.0-19.3) | 16.4 (1.5-26.6) | 9.0 (0.9-15.3) | 10.6 (3.5-22.4) | 5.8 (1.9-13.4) |
| Si_IO | 16.2 (4.8-27.5) | 11.0 (3.2-19.0) | 13.2 (5.6-21.7) | 9.1 (4.0-15.1) | 11.2 (0.5-17.9) | 8.0 (0.4-11.4) |
| Si_VL | 26.0 (7.5-39.6) | 16.2 (4.4-24.2) | 19.2 (4.1-33.1) | 11.1 (2.5-19.8) | 15.0 (0.6-30.1) | 8.6 (0.3-18.4) |
| Si_IO_VL | 11.8 (1.0-20.4) | 8.3 (1.1-16.0) | 9.6 (3.4-18.5) | 6.8 (2.4-13.1) | 7.7 (1.8-15.5) | 5.7 (1.2-10.8) |
| St_Ba | 24.7 (6.8-35.0) | 11.5 (3.1-16.2) | 15.8 (2.0-26.0) | 7.5 (0.9-12.3) | 11.9 (2.9-22.0) | 5.7 (1.2-11.1) |
| St_IO | 17.8 (4.7-25.7) | 8.8 (2.2-12.8) | 10.2 (2.3-17.8) | 5.1 (1.1-8.7) | 8.1 (2.0-16.5) | 4.0 (1.2-8.5) |
| St_RG | 8.5 (3.2-16.3) | 4.5 (2.0-8.4) | 5.9 (1.5-11.8) | 3.2 (0.8-5.8) | 3.4 (0.7-7.4) | 1.8 (0.4-4.0) |
| St_IO_RG | 11.9 (3.9-18.0) | 7.1 (2.5-11.4) | 7.2 (1.3-12.2) | 4.5 (0.8-7.1) | 4.8 (0.9-9.9) | 2.9 (0.7-6.0) |

The largest variability among predictions is observed for the Si_IO_VL model, while the St_IO_RG model has the lowest variability.

### Prediction of the impact of behavioural interventions

Figures 2–5 show that the interventions have a similar effect on HIV relative to the Simpact Cyan 1.0 and StepSyn 1.0 basic models (Si_Ba and St_Ba).

**Figure 2.**
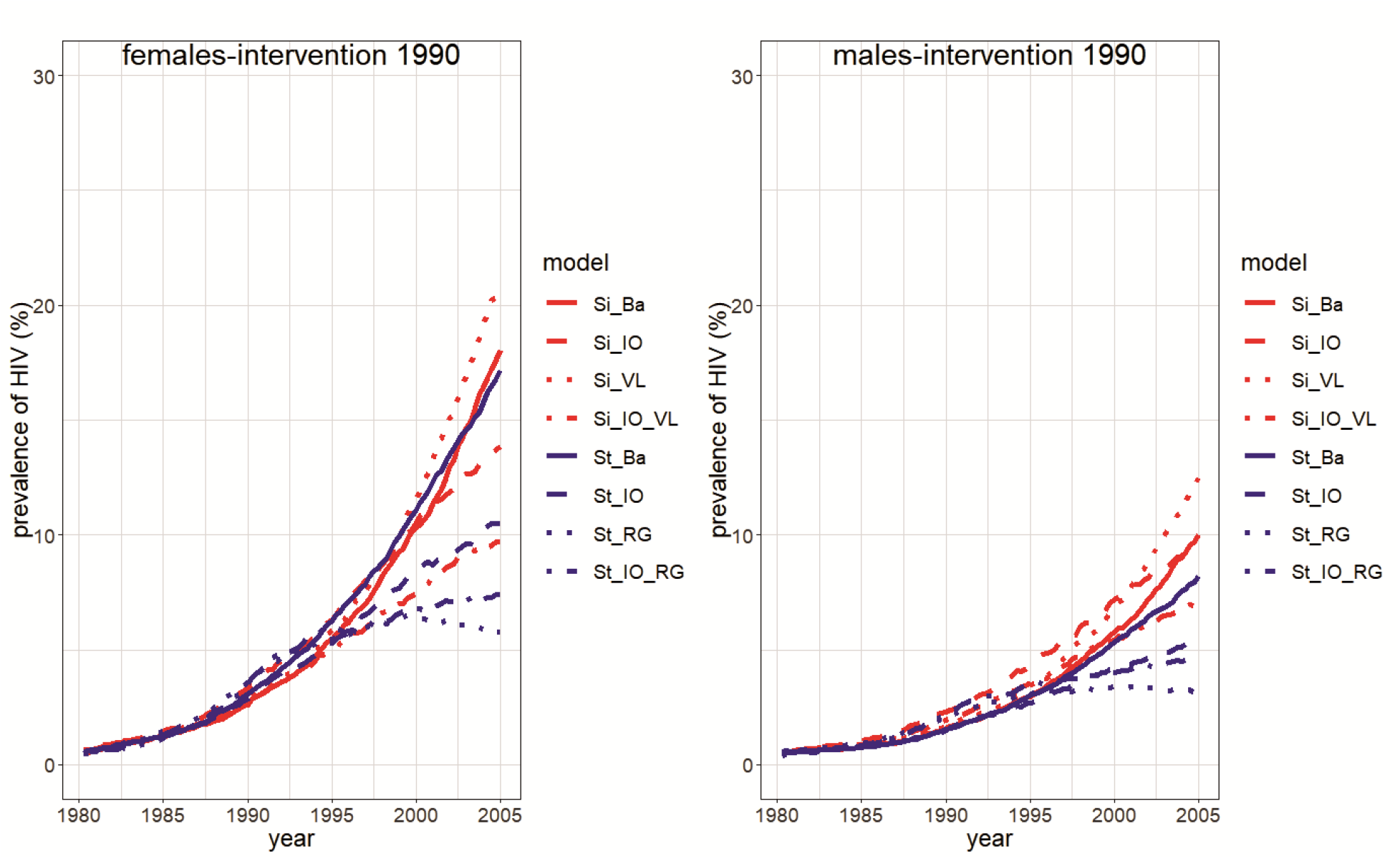
Prevalence curves for HIV from 1980 to 2005 in case a behavioural intervention is implemented in 1990. Median HIV prevalence (in %) of 100 simulations. Left: females; right: males. Models: Si_Ba: Simpact 1.0 basic model; Si_IO: Simpact 1.0 model with inflow and outflow; Si_VL: Simpact 1.0 model with VL-dependent HIV transmission hazard; Si_IO_VL: Simpact 1.0 model with inflow, outflow and VL-dependent HIV transmission hazard; St_Ba: StepSyn 1.0 basic model; St_IO: Stepsyn 1.0 model with inflow and outflow; St_RG: StepSyn 1.0 model with STI life history explicitly modeled; St_IO_RG: StepSyn 1.0 model with inflow, outflow and STI life history explicitly modeled.

Apart from reducing HIV, the interventions also reduce HSV-2 (see Supplementary Table S25), and the size of the reduction differs among the eight models and between interventions (intervention in 1990: reduction of the median HSV-2 prevalence in 1997: 0.3%-3.6% (females) and 0.2%-2% (males); lower promiscuity in 1980: reduction of 0.7%-5.3% (females) and 0.7%-2.9% (males)).

Median HIV prevalence in 2004 for females and males (in %) and range (the maximum minus the minimum) of 100 simulations in case of no intervention, a behavioural intervention in 1990 and lower promiscuity from 1980 onwards are shown in Table 1. For all models, the implementation of a behavioural intervention reduced the median HIV prevalence and the range (Table 1 and Fig. 4–5). After implementing a behavioural intervention in 1990, a reduced but still increasing HIV epidemic was observed for all models except the StepSyn 1.0 model with STI life history explicitly modeled (St_RG) which shows a decreased epidemic after 1998 (Fig. 2). In case of lower promiscuity from 1980 onwards (Fig. 3), similar trends are observed but the reduction in HIV prevalence is considerably larger than when applying the intervention in 1990. For the two models that resulted in the best predictions of the HIV prevalence in 2004 in case of no intervention (Si_IO_VL and St_IO_RG), the predicted effect of a behavioural intervention was larger for the St_IO_RG model than for the Si_IO_VL model (Table 1). While the Si_IO_VL model predicts a 2.2% and 1.5% reduction in median HIV prevalence in 2004 for females and males respectively, in case a behavioural intervention was implemented in 1990, the St_IO_RG model predicts a 4.7% and 2.6% reduction for females and males respectively. In case of lower promiscuity from the start of the epidemic (1980), Si_IO_VL predicts a 4.1% and 2.6% reduction in HIV prevalence in 2004 for females and males respectively, while St_IO_RG model predicts a 7.1% and 4.2% reduction in females and males respectively.

**Figure 3.**
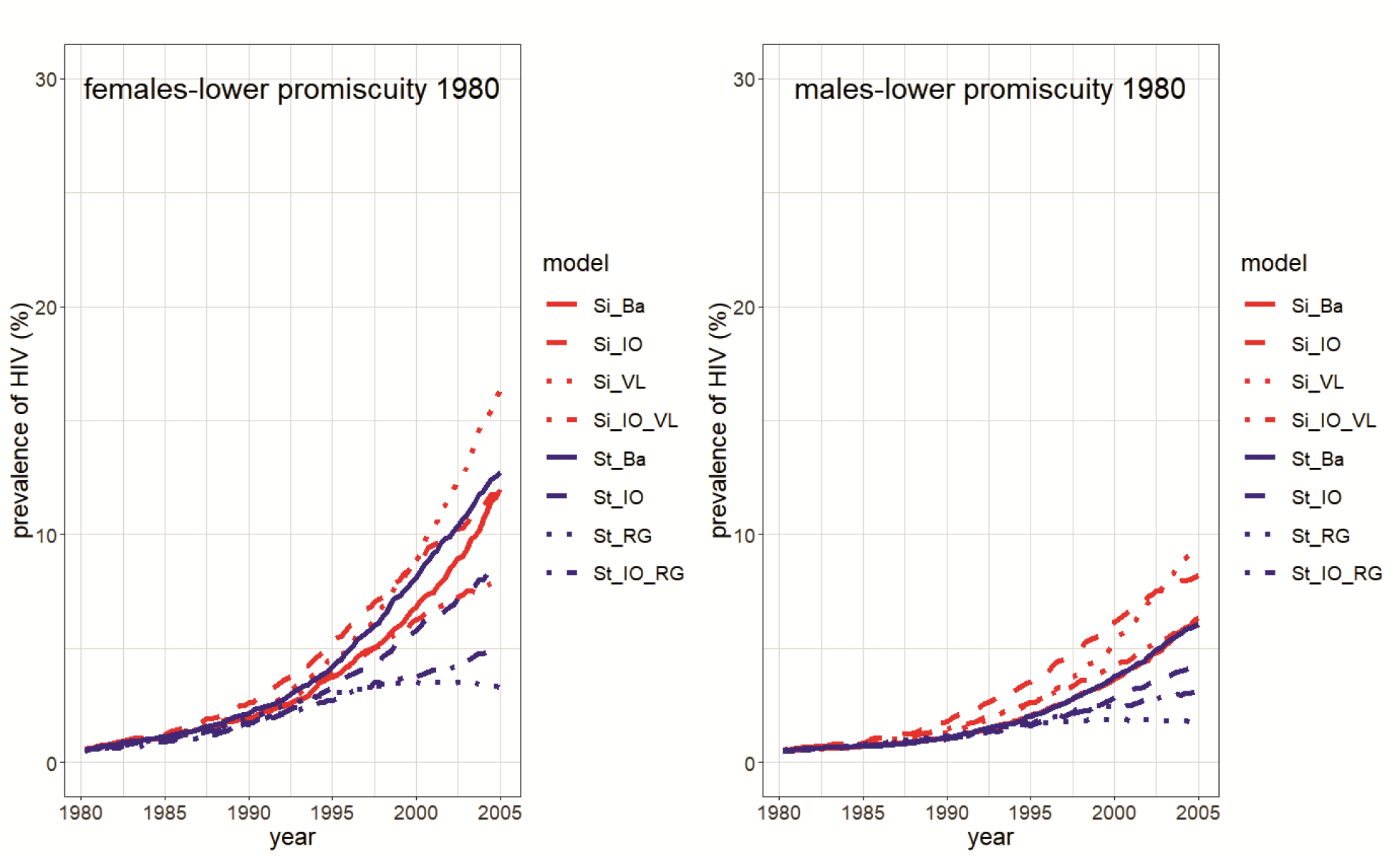
Prevalence curves for HIV from 1980 to 2005 in case of lower promiscuity from 1980 onwards. Median HIV prevalence (in %) of 100 simulations. Left: females; right: males. Models: Si_Ba: Simpact 1.0 basic model; Si_IO: Simpact 1.0 model with inflow and outflow; Si_VL: Simpact 1.0 model with VL-dependent HIV transmission hazard; Si_IO_VL: Simpact 1.0 model with inflow, outflow, VL-dependent HIV transmission hazard; St_Ba: StepSyn 1.0 basic model; St_IO: Stepsyn 1.0 model with inflow and outflow; St_RG: StepSyn 1.0 model with STI life history explicitly modeled; St_IO_RG: StepSyn 1.0 model with inflow, outflow and STI life history explicitly modeled.

**Figure 4.**
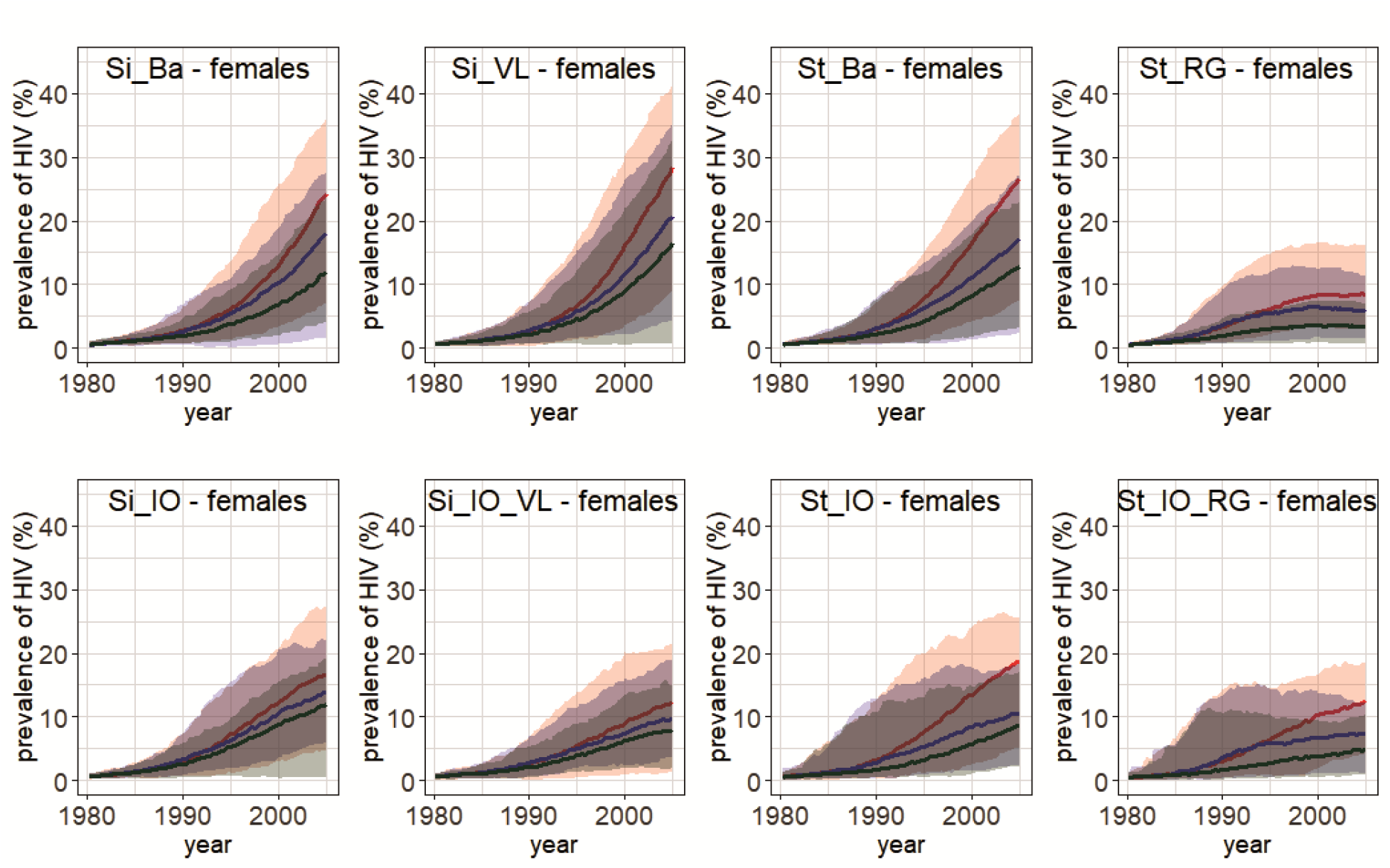
Prevalence curves for HIV in females from 1980 to 2005. Median HIV prevalence (in %) of 100 simulations and the range ([minimum,maximum]) (shaded area). Red: no intervention implemented; blue: intervention implemented in 1990; green: lower promiscuity from 1980 onwards. Models: Si_Ba: Simpact 1.0 basic model; Si_IO: Simpact 1.0 model with inflow and outflow; Si_VL: Simpact 1.0 model with VL-dependent HIV transmission hazard; Si_IO_VL: Simpact 1.0 model with inflow, outflow and VL-dependent HIV transmission hazard; St_Ba: StepSyn 1.0 basic model; St_IO: Stepsyn 1.0 model with inflow and outflow; St_RG: StepSyn 1.0 model with STI life history explicitly modeled; St_IO_RG: StepSyn 1.0 model with inflow, outflow and STI life history explicitly modeled.

**Figure 5.**
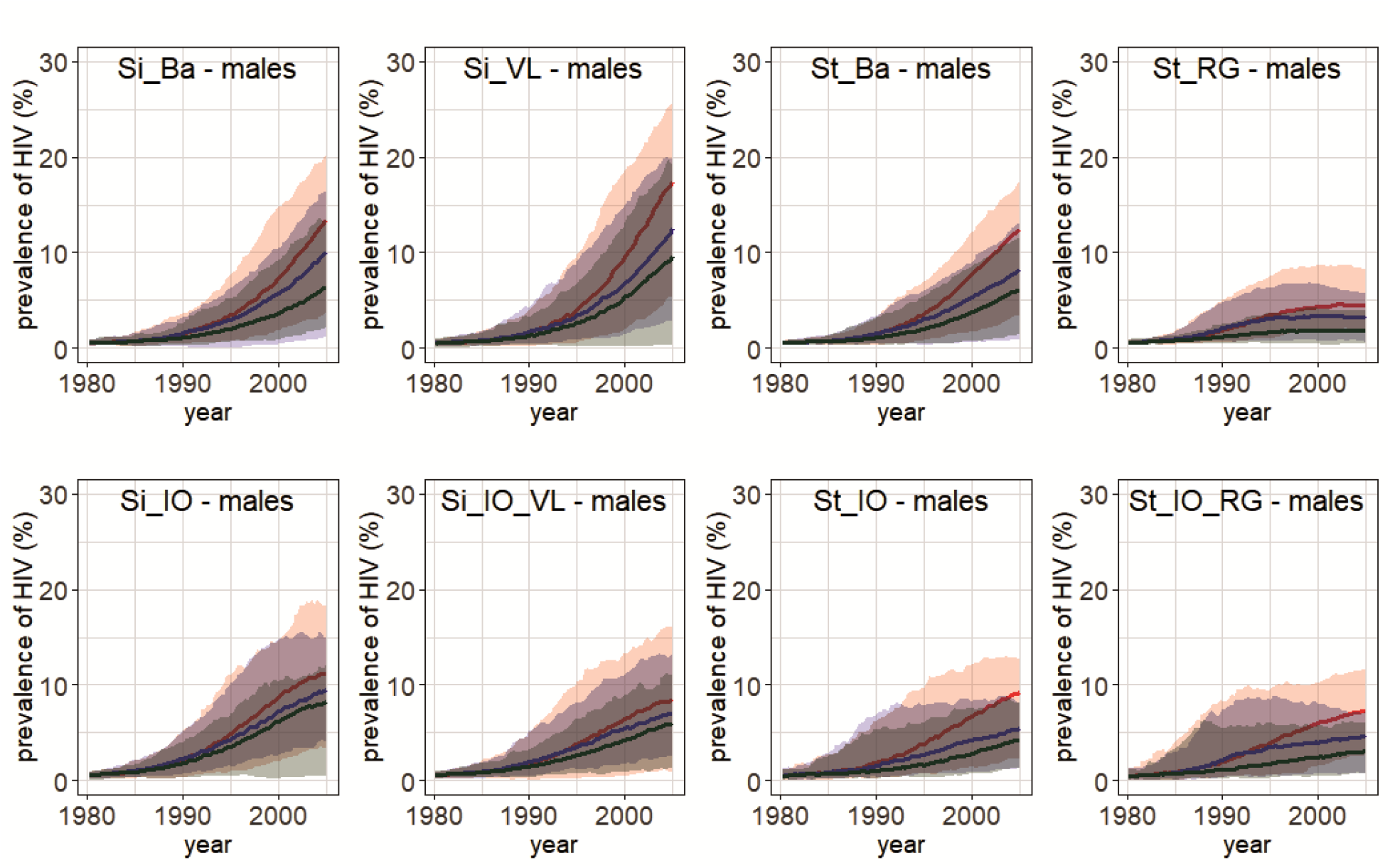
Prevalence curves for HIV in males from 1980 to 2005. Median HIV prevalence (in %) of 100 simulations and the range ([minimum,maximum]) (shaded area). Red: no intervention implemented; blue: intervention implemented in 1990; green: lower promiscuity from 1980 onwards. Models: Si_Ba: Simpact 1.0 basic model; Si_IO: Simpact 1.0 model with inflow and outflow; Si_VL: Simpact 1.0 model with VL-dependent HIV transmission hazard; Si_IO_VL: Simpact 1.0 model with inflow, outflow, VL-dependent HIV transmission hazard; St_Ba: StepSyn 1.0 basic model; St_IO: Stepsyn 1.0 model with inflow and outflow; St_RG: StepSyn 1.0 model with STI life history explicitly modeled; St_IO_RG: StepSyn 1.0 model with inflow, outflow and STI life history explicitly modeled.

## Discussion

In this study, eight individual-based models, generated with the simulators Simpact Cyan 1.0 and StepSyn 1.0 were compared in terms of their prediction of the impact of behavioural interventions on the course of the HIV epidemic in Yaoundé, Cameroon.

All models could be fitted equally well to the calibration targets in Supplementary Table S3 (see Supplementary Figure S12), which shows that similar HIV prevalence curves can be simulated using different model assumptions and transmission parameters.

After calibration of the models, they were first validated against data not used for fitting. For each of the two modelling frameworks, the model that implements population inflow and outflow together with a detailed description of HIV transmission (i.e. the Si_IO_VL and St_IO_RG models) shows the best prediction of the HIV prevalence for males and females in 2004 (Fig. 1).

In general, a model implementing inflow and outflow better predicts HIV prevalence at a time point not used for fitting than its counterpart without inflow and outflow (Fig. 1). Remarkably, while models with no inflow and outflow implementing a generic STI co-factor effect largely overestimated the HIV prevalence in 2004, the model implementing HSV-2 life history and its effect on HIV in detail, and no inflow and outflow (the St_RG model) predicted a lower HIV prevalence for both males and females than what was reported in the literature. Moreover, the effect of implementing inflow and outflow on the model predictions was smaller for the model implementing HSV-2 in detail than for the other models.

The predictions of the impact of behavioural interventions for the eight models are similar during the first five years after the intervention, but show large differences on the long term (see Fig. 2 and Fig. 3). Similar conclusions could be drawn by Eaton et al.^23^ when comparing mathematical models predicting the impact of antiretroviral therapy (ART).

All simplified Simpact Cyan 1.0 models (Si_Ba, Si_IO, Si_VL) overestimate the HIV prevalence in 2004 compared to the most detailed model (Si_IO_VL). As a consequence, they also predict larger effects of interventions (Figures 4–5, Table 1, Supplementary Figures S14-S15). For StepSyn 1.0, the models that overestimate (St_Ba, St_IO) and underestimate (St_RG) the HIV prevalence, predict larger and smaller effects, respectively, compared to the most detailed model (St_IO_RG) (Figures 4–5, Table 1, Supplementary Figures S14-S15).

As the Si_IO_VL and St_IO_RG models showed the best predictions of the HIV prevalence in 2004, these models can be considered as the most reliable for predicting behavioural interventions.

Although the Si_IO_VL and St_IO_RG models predict the HIV prevalence in 2004 equally well, the two models behave differently when implementing the effect of behavioural intervention. After an intervention in 1990, the median HIV prevalences for females and males in 2004 were reduced with 2.2% and 1.5% respectively for the Si_IO_VL model (see Table 1), while the St_IO_RG model predicts a reduction of 4.7% for females and 2.6% for males. In case of lower promiscuity from 1980 onwards, a reduction in HIV prevalence of 4.1% for females and 2.6% for males were predicted by the Si_IO_VL model. For the St_IO_RG model, the corresponding values were 7.1% and 4.2%. More data has to become available to further explore which of these two models provides the most reliable predictions of behavioural interventions.

Seven of the eight models predicted that the HIV epidemic would still increase during the period 1980-2005, although at a lower rate, after applying a behavioural intervention (see Fig. 2 and Fig. 3). Only the St_RG model predicted that the HIV epidemic will decrease after reaching a peak in 1997.

This study shows that differences in model assumptions and model complexity can considerably influence their predictions of the impact of behavioural interventions. Hontelez et al.^24^ reported similar conclusions after stepwise inclusion of model complexity in a model for predicting the impact of a universal test and treat (UTT) intervention and concluded that sufficient detail is necessary to make accurate predictions.

Apart from differences in the size of the impact of a behavioural intervention on the HIV epidemic, we also detected differences in qualitative behaviour between simulations generated with different models.

Furthermore, we showed that even when there is an agreement between two models in their prediction of a future time point not used for fitting, they can have different outputs when simulating the impact of interventions. Without more HIV prevalence data for validation, it is not possible to determine which of these two models is the most reliable. In case more HIV prevalence data would have been available for the period 1999-2005 than only the HIV prevalence in 2004, validation could have been performed on multiple data points not used for calibration. This would have enabled to better determine which model has the best prediction of the epidemic for the period 1999-2005. These findings highlight the importance of making more data available for both the calibration and validation of epidemiological models that aim to inform decisions made by policy-makers.

## Acknowledgements

The research conducted by D.M.H. and N.H. in this study was funded by the Fonds Wetenschappelijk Onderzoek - Vlaanderen (Research Foundation – Flanders; FWO, http://www.fwo.be/en/) (Grant agreements G0E8416N and G0B2317N). The research done by J.D.S. and A.M.V. in this study has been supported in part by grants G.0692.14 and G0B2317N, funded by the FWO, Belgium. P.L. was supported by a PhD grant of the FWO (1S31916N). WD was supported by a postdoctoral fellowship from FWO (12L5816N). Research done by V.M. in this study has been funded by the ELTE Institutional Excellence Program supported by the National Research, Development and Innovation Office (NKFIH-1157-8/2019-DT). V.M. was also supported by the ÚNKP-18-4 New National Excellence Program of the Hungarian Ministry of Human Capacities and by a Bolyai János Research Fellowship of the Hungarian Academy of Sciences. The authors gratefully acknowledge support from the FWO Scientific Research Community on Network Statistics for Sexually Transmitted Diseases Epidemiology. The computational resources and services used in this work were provided by the VSC (Flemish Supercomputer Center), funded by the Research Foundation - Flanders (FWO) and the Flemish Government – department EWI.

## Author contributions

All authors have participated in the research and article preparation. D.M.H. wrote the R code for calibrating the models and generating the figures in this paper with support of N.H. and W.D., and performed the model comparison with support of J.D.S. J.L., W.D. and N.H. developed SimpactCyan 1.0. J.D.S. developed StepSyn 1.0 with support of P.J.K.L., V.M. and A.M.V. J.D.S and P.J.K.L. fitted power laws to behavioural data to generate the sexual network for StepSyn 1.0. All authors contributed to writing this manuscript. All authors have approved the final article.

## Data availability

The data used to calibrate the models in this study, together with references supporting these data, are available within the Supplementary Material, section 2.

The R-scripts that have been used for fitting HIV transmission parameters are available from GitHub: https://github.com/dmhendrickx/Scripts_comparison_Simpact_StepSyn

## Supplementary Material

see Supplementary_material.pdf

