## Supplementary Material for "Comparison of two individual-based model simulators for HIV epidemiology in a population with HSV-2 using as case study Yaoundé-Cameroon, 1980-2005"

[1]I-BioStat, Data Science Institute, Hasselt University, Hasselt, Belgium
[2]KU Leuven – University of Leuven, Department of Microbiology and Immunology, Rega Institute for Medical Research, Clinical and Epidemiological Virology, B-3000, Leuven, Belgium
[3]Global Health and Tropical Medicine (GHTM), Unidade de Microbiologia Médica, Instituto de Higiene e Medicina Tropical, Universidade Nova de Lisboa, Lisbon, Portugal
[4] Artificial Intelligence Lab, Department of Computer Science, Vrije Universiteit Brussel, 1050 Brussels, Belgium
[5] The South African Department of Science and Technology-National Research Foundation (DST-NRF) Centre of Excellence in Epidemiological Modelling and Analysis (SACEMA), Stellenbosch University, Stellenbosch, South Africa
[6]Department of Global Health, Faculty of Medicine and Health, Stellenbosch University, Stellenbosch, South Africa
[7]International Centre for Reproductive Health, Ghent University, Ghent, Belgium
[8]School for Data Science and Computational Thinking, Stellenbosch University, Stellenbosch, South Africa
[9]Expertise Centre for Digital Media, Hasselt University – tUL, Diepenbeek, Belgium
[10]Centre for Health Economics Research and Modelling Infectious Diseases, Vaccine & Infectious Disease Institute, University of Antwerp, Antwerp, Belgium
[11]Institute of Biology, Eötvös Loránd University, Budapest, Hungary
[*]
[+]these authors contributed equally to this work

#### 1. Key model differences between Simpact 1.0 and StepSyn 1.0

The Simpact Cyan 1.0 and StepSyn 1.0 modelling frameworks were developed separately. Similarities and differences between the two modelling frameworks are outlined in Table S1. Models developed with Simpact Cyan 1.0 and StepSyn 1.0 are both individual-based and both include HIV natural history. While StepSyn models can also include the natural history of other STIs, Simpact Cyan 1.0 models are restricted to a generic, constant STI co-factor effect on HIV. Within the StepSyn 1.0 framework, five different STIs (other than HIV) can be simulated but for the present study, only HSV-2 and HIV were simulated.

Models developed with Simpact Cyan 1.0 and StepSyn 1.0 are both accessible from R, but they differ in the way they are implemented. While the source code for StepSyn is R code, Simpact Cyan 1.0 is implemented in C++ with interfaces to Python and R. In this way, Simpact Cyan 1.0 combines the computational efficiency of C++ with the user-friendliness of R and Python.

Simpact Cyan 1.0 and StepSyn 1.0 also differ in the way the state of the model system is updated and, as a consequence, in the way stochastic processes are described. StepSyn implements discrete time steps of one week and updates population, sexual links, and STI states at each time step. Stochastic processes are implemented through R probability distribution functions which are called in each time step. Simpact Cyan 1.0 implements a continuous-time model and the state of the model system is updated each time an event happens, and therefore requires the use of hazard functions. The hazard at a certain time point is defined as the event rate at that particular time point, conditional on model states such as still being alive at that time point.

An advantage of the continuous-time implementation is that events that happen on different time scales can be integrated into a single simulation. A disadvantage is that continuous-time models are computationally more complex, and therefore require a longer simulation time.

Another difference between both modelling frameworks is the way individuals can enter and leave the population. In Simpact Cyan 1.0, individuals can enter the population by birth, and leave the population by non-AIDS or AIDS mortality. The mortality events are age-dependent. StepSyn 1.0 is not age-structured; individuals enter the population by immigration or sexual debut and leave the population by emigration, AIDS- or non-AIDS age-independent mortality.

An extra option in Simpact Cyan 1.0 that is not implemented in StepSyn 1.0 is pregnancy. In Simpact Cyan 1.0, women are pregnant between a conception event and a birth event. This means that after a simulation run, one can measure how many women were pregnant at any given point in time in the simulation.

Formation and break-up of relationships in the sexual network are implemented differently in StepSyn 1.0 and Simpact Cyan 1.0. In Simpact Cyan 1.0, formation and break-up of relationships happen through formation and dissolution events. The timing of these events is sampled from a probability distribution that emerges as a result of the hazard function that was specified. In StepSyn, formation and break-up of relationships is performed each time step, using an individual-specific probability of forming a new relationship.

An extra option in Simpact Cyan 1.0 that is not currently implemented in StepSyn 1.0 is the simulation of HIV diagnosis and treatment, including treatment dropout, and the possibility to introduce treatment during the simulation through a so-called intervention event.

#### 1.1 Simpact Cyan 1.0 sexual network

In Simpact Cyan 1.0, formation and dissolution of relationships in the sexual network are simulated as discrete events. For man-woman pairs of sexually active persons, formation events are scheduled. When a formation event is triggered, a sexual relationship is established, and a dissolution event is scheduled. When the dissolution event is triggered, the relationship ceases to exist.

Formation and dissolution events are described by hazards. In the description of the hazards, parameters can be included describing the dependence of relationship formation and dissolution on the number of relationships the person already has, the age of the person and the preferred age gap between the partners.

In this study, the formation hazard is described by the following formula:

$$hazard = F \times exp(\alpha_{baseline} + \alpha_{numrel,man} P_{man} + \alpha_{numrel,woman} P_{woman}) \quad (1)$$

where $P_{man}$ and $P_{woman}$ are the number of partners the man and the woman already have, respectively.

The parameter $\alpha_{baseline}$ represents a baseline value for relationship formation (0.1 by default). The parameters $\alpha_{numrel,man}$ and $\alpha_{numrel,woman}$ represent the influence of the number of partners the man and the woman already have, respectively (0 by default). Decreasing $\alpha_{numrel,man}$ and $\alpha_{numrel,woman}$ decreases the formation hazard. Therefore these parameters are used to describe the behavioural interventions in this study (see section 6 of this Supplementary Material).

To avoid that the number of relationships automatically increases with the population size, a normalization factor F is applied, which roughly divides the hazard by the size of the population.

The dissolution hazard in this study is described by a baseline value $\alpha_0$ for relationship dissolution (0.1 by default): $hazard = \exp(\alpha_0)$.

In this comparison study, the default settings of Simpact Cyan 1.0 for the formation and dissolution hazard were applied for the scenarios without behavioural intervention (the developers of Simpact Cyan 1.0 did not have access to the behavioural data of the city of Yaoundé described in paragraph 1.2.1).

Because Simpact Cyan 1.0 keeps track of the history of each individual, we can reconstruct the sexual network from the model output. The network can be reconstructed at a given time point by including all relationships a person has at that time point for each person in the population. Furthermore, a cumulative network over a certain period of interest can be reconstructed, including all relationships between individuals in the population during that period (e.g. all relationships people had during the past 12 months).

#### 1.2. StepSyn 1.0 sexual network

##### *1.2.1. Data available for generating the sexual network in StepSyn 1.0*

Since StepSyn 1.0 models explore behavioural differences between individuals, behavioural data on the distribution of the number of non-marital partners in the last 12 months were gathered for Yaoundé during the development of the StepSyn 1.0 modelling framework. These data were obtained from the surveys of the 4 Cities Study and provided by the Institute of Tropical Medicine (ITM) in Antwerp, Belgium (Anne Buvé, unpublished database). These data were used to estimate the parameters of a power-law distribution to generate the sexual network for the four StepSyn 1.0 models (see paragraph "Models used in this study" in the Methods section of the main text). This work was performed before the current study.

##### *1.2.2. Generation of the sexual network*

Power law distributions (separately for men and women) were fitted to the data on the reported number of partners in the last 12 months taken from the 4 Cities Study. Each individual is assigned a preferred degree (number of partners within 12 months) drawn from the appropriate distribution. The individual-specific preferred degrees (PDi) are translated into probabilities of forming a new short term relationship per week (PFi), which are governed by the shortage of short links to reach the preferred degree and calculated using the formula PFi = PDi / (MD + 52), with MD being the mean duration of short term relationships in weeks. With these parameters, in each week, the total male demand for new relationships is higher than the female one, partly because of the male-biased sex ratio, and partly because of female underreporting of the number of partners in the survey that serves as the basis for the distribution used. The extra male demand is uniformly distributed between the females, a method that is acceptable for modelling purposes in the absence of data on mixing patterns and life-cycle changes in sexual behaviour [1]. Although the assumption of uniformity can lead to heterogeneity in female activity being understated, assuming that the extra male demand is distributed only to the more active women would over-estimate the number of partners of females with high activity, while leaving the number of partners of females with low activity unchanged [1]. The number of weekly sex acts in married couples and short-term relationships is calculated by applying two different Poisson distributions so that their means would be consistent with the values found in the 4 Cities Study for Yaoundé [2,3].

The parameter for the proportion of male pending short links is equal to 1 by default, which means that all male pending short links are fulfilled. Lowering this parameter decreases the probability that relationships are formed. Therefore, this parameter was used to describe the behavioural interventions (see section 6 of this Supplementary Material).

##### 1.3. Simpact Cyan 1.0 HIV natural history

In Simpact Cyan 1.0, an infected person will go through the following HIV stages [7]:

- acute HIV infection of 12 weeks [19] (parameter in Simpact Cyan 1.0: chronicstage.acutestage);
- chronic HIV infection – until 1 year before dying of AIDS [20,21] (parameter in Simpact Cyan 1.0: agestage.start);
- AIDS stage: 1 year before dying of AIDS until 6 months before dying of AIDS (default);
- final AIDS stage, the person is too ill to be sexually active: 6 months before dying of AIDS until AIDS-related death (default).

In general, the formula for survival time in Simpact Cyan 1.0 is [7]:

$$t_{survival} = \frac{C}{V_{sp}^{-k}} \times 10^{X}$$

Where $V_{sp}$ is a person's set point viral load. In this study x = 0 (default), k = 0 (parameter in Simpact Cyan 1.0: mortality.aids.survtime.k) and C = 2 (parameter in Simpact Cyan 1.0: mortality.aids.survtime.C). The time to AIDS-related death is implemented as the time of infection plus the survival time (here: 2 years, C = 2).

##### 1.4. StepSyn 1.0 HIV natural history

In StepSyn 1.0, the duration of the acute HIV stage in this study is 12 weeks [19] (parameter names in StepSyn 1.0: hiv.strain1.acute.dur.min and hiv.strain1.acute.dur.max). The minimum time to develop AIDS after being HIV infected is 2 years [20,22] (parameter name in StepSyn 1.0: hiv.strain1.aids.progr.start.time). The mean time to develop AIDS is 9 years [20,22] (parameter in StepSyn 1.0: hiv.strain1.aids.progr.mean.time). The time to death when having AIDS is 1 year [20,22] (parameter in StepSyn 1.0: hiv.strain1.aids.dur).

**Table S1.** Comparison of properties of models developed with Simpact Cyan 1.0 and StepSyn 1.0.

| Model property | Simpact Cyan 1.0 | StepSyn 1.0 |
|---|---|---|
| Simulation framework | Individual-based model | Individual-based model |
| Implementation language | C++ with R and Python interfaces | R |
| Source code freely available | Yes | After the publication of the manuscript describing the modelling framework in detail |
| Update of the state of the model system | Event-driven: the state of the model system is updated each time an event happens | Fixed time steps of one week, with the creation and dissolution of sexual relationships and STI and HIV transmission |
| Description of stochastic processes | By hazard functions | By probability distribution functions called in each time step |
| Age-structured | Yes | No |
| Entering the population | Birth (birth event) | Sexual debut (birth-rate), immigration |
| Leaving the population | AIDS- and non-AIDS mortality (mortality event, age-dependent) | AIDS- and non-AIDS mortality (mortality rate), emigration |
| Formation and break-up of relationships within the sexual network | Formation and dissolution event. Timing of events sampled from a probability distribution resulting from carefully chosen hazard functions. | Each time step, using an individual-specific probability of forming a new relationship |
| Individual variation in sexual behaviour | In the formation hazard, the eagerness of a person to form a relationship and the preferred age gap are used, to allow for individual variation in sexual behaviour. Both values are drawn from user-specified probability distributions. | Number of short term relationships follows a power-law distribution |
| HIV natural history / stages | Yes (acute phase, chronic phase, AIDS stage) | Yes (acute phase, chronic phase, AIDS stage) |
| Other STIs as co-factor for HIV infection | Yes (implemented as a simplified STI transmission hazard) | Yes (syphilis, chancroid, gonorrhoea, chlamydia, HSV-2) |
| Natural history/stages – other STIs | No, stages are not explicitly modelled | Yes, with genital ulcers and discharge explicitly modelled |
| Pregnancy | Yes, as the period between a conception event and a birth event. | No |
| Diagnosis / treatment of HIV | Yes | No |
| HIV seeding | A fraction to specify the probability of each person in the group to be a seeder or an amount to specify the number of seeders. Fraction or amount is taken from the number of people in the population. | Either a random binomial for HIV seeding (seeding varies between simulations) or a fixed fraction to specify the fraction of seeders; this fraction is taken from the number of males and the number of females separately. |
| Interventions | Behavioural interventions, HIV treatment | Behavioural interventions, circumcision, treatment of syphilis, chancroid, gonorrhoea, chlamydia, HSV-2 |
| individual variation of viral load | Implemented in HIV transmission hazard as hazard | No |
| Computational time for 10,000 simulations* | ~35 hours | ~5 hours |

*on 100 cores of the VSC (Xeon E5-2680v2 CPUs 2.8 GHz, 25 MB level 3 cache). Population size ~ 9000; time frame of the simulation = 35 years; parameters: see Table S13 (Simpact Cyan 1.0 basic model) and S17 (StepSyn 1.0 basic model).

#### 2. Data used for fitting Simpact Cyan 1.0 and StepSyn 1.0 models

For the comparison of the two models, demographic data for 1997, and HIV prevalence data for 1989–1998 related to the African city of Yaoundé (Cameroon), are used. The adult male population (15-59 years) for that year was estimated at 387,398, in the 4 Cities Study [4]. Based on the surveys made by the same study (age range 15-49 years), 34.5% of men and 44.2% of women were married; and 7.2% of married men were polygamous [3]. Assuming most of the latter had 2 wives, we estimate the number of women as 387,398 ∗ 0.345 ∗ (1+0.072) / 0.442 = 324,152, implying an adult sex ratio (M:F) of 1000:836.74.

Since one of our aims here is to compare HIV prevalence curves resulting from fitting HIV transmission parameters of our two models to data, we gathered HIV-1 prevalence data for Yaoundé for the period 1989–1998. The only prevalence data available were of pregnant women, for whom prevalence increased from 0.7% in 1989 to 5.5% in 1998 (US Census Bureau, 2001, HIV/AIDS Profile, Cameroon, HIV/AIDS Surveillance Database [5]) as shown in Table S2.

**Table S2.** HIV-1 prevalence data for Yaoundé's pregnant women. Source: US Census Bureau, 2001, HIV/AIDS Profile, Cameroon, HIV/AIDS Surveillance Database [5] (confidence interval and sample size not available).

| Year | HIV-1 prevalence (%) |
|---|---|
| 1989 | 0.71 |
| 1990 | 1.32 |
| 1991 | 2.10 |
| 1992 | 1.91 |
| 1993 | 1.30 |
| 1994 | 3.00 |
| 1995 | 2.72 |
| 1996 | 4.81 |
| 1998 | 5.51 |

Because StepSyn 1.0 does not implement pregnant women as a separate category of the population, and we wanted to compare model simulations for males and females separately, the prevalence rates for pregnant women were converted to prevalence rates for males and females as follows. In 1997, during the 4 Cities Study, HIV-1 prevalence was 4.1% (95% confidence interval: CI: 3.0% - 5.7%; sample size: n = 896) in men (age 15-49 years) and 7.8% (CI: 6.2% - 9.6%; n = 1017) in women (age 15-49 years) [6]. The prevalence for pregnant women in 1997 was estimated by fitting a smoothing spline through the data from Table S2, using the smooth.spline function of the stats package in R, and evaluating the spline at the time point corresponding with 1997 (see Figure S1). We obtained a prevalence of 4.778 % for pregnant women in 1997 and calculated the prevalence ratio of men/pregnant women (resp. women/pregnant women) by dividing 4.1 (resp. 7.8) by 4.778. The prevalence ratios (0.858 for men and 1.632 for women) were multiplied with the data from Table S2 to obtain HIV prevalence rates for men and women separately for all years between 1989 and 1998 (see Table S3).

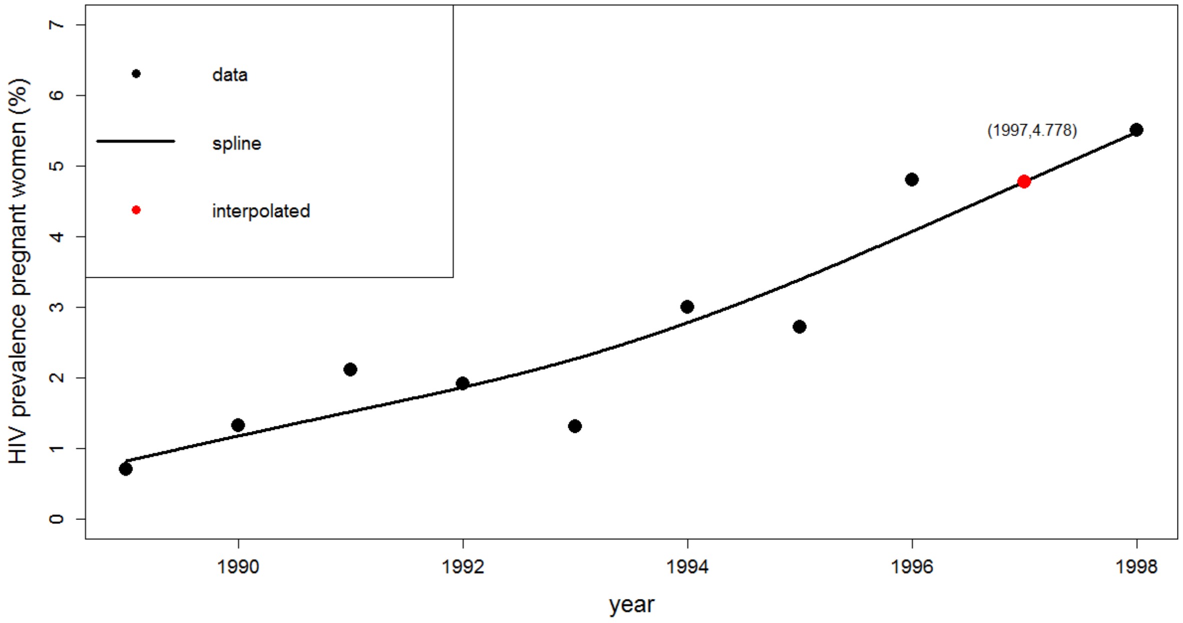


**Figure S1.** Interpolation spline through the data of Table S2 to obtain the HIV prevalence in 1997 for pregnant women. The smoothing parameter was determined using Generalized Cross-Validation (GCV).

**Table S3.** Estimated HIV-1 prevalence rates for Yaoundé's men and women, that were used as model calibration targets in this study.

| Year | HIV-1 prev (%) men | HIV-1 prev (%) women |
|---|---|---|
| 1989 | 0.609 | 1.159 |
| 1990 | 1.133 | 2.154 |
| 1991 | 1.802 | 3.427 |
| 1992 | 1.639 | 3.117 |
| 1993 | 1.115 | 2.122 |
| 1994 | 2.574 | 4.896 |
| 1995 | 2.334 | 4.439 |
| 1996 | 4.127 | 7.850 |
| 1997 | 4.100 | 7.800 |
| 1998 | 4.728 | 8.992 |

### 3. Calibration of the models

#### 3.1. Overview of Simpact Cyan 1.0 parameters that were fitted to calibration targets

Table S4 gives an overview of the Simpact Cyan 1.0 parameters that were fitted to the calibration targets in Table S3, together with the initial range ([minimum, maximum]) used for fitting.

**Table S4.** Overview of the Simpact Cyan 1.0 parameters that were fitted to the calibration targets in Table S3. The last column indicates in which models these parameters were used (see main text for abbreviations).

| parameter definition | parameter name | initial range | models |
|---|---|---|---|
| HIV baseline transmission hazard a<br>(annotated by hiv_a in the figures) | hivtransmission.param.a | [-3.65, -1.34] | all |
| HIV transmission hazard - influence of the HIV-infected person (index) being infected with HSV-2<br>(annotated by hiv_e1 in the figures) | hivtransmission.param.e1 | [0, 1.4] | all |
| HIV transmission hazard - influence of the HIV-uninfected person (exposed) being infected with HSV-2<br>(annotated by hiv_e2 in the figures) | hivtransmission.param.e2 | [0, 1.4] | all |
| HIV transmission hazard – influence of gender<br>(annotated by hiv_f1 in the figures) | hivtransmission.param.f1 | [0.7, 1.4] | all |
| HIV transmission hazard – parameter b in the formula<br>$hazard = \exp(a + bV^{-c} + other\ terms)$ [7]<br>where<br>a = baseline transmission hazard (see above)<br>V = current viral load | hivtransmission.param.b | [-15,-10] | Si_VL<br>Si_IO_VL |
| HIV transmission hazard – parameter c in the formula<br>$hazard = \exp(a + bV^{-c} + other\ terms)$ [7]<br>where<br>a = baseline transmission hazard (see above)<br>V = current viral load | hivtransmission.param.c | [0.1, 0.2] | Si_VL<br>Si_IO_VL |

In this study, the HIV transmission hazard in the most detailed model (Si_IO_VL) is given by the following formula :

$$hazard = \exp(a + bV^{-c} + Wf_1 + e_1 HSV2_{infected} + e_2 HSV2_{uninfected})$$

In this formula V is the current viral load of the HIV infected person; W is a binary factor which is 1 if the uninfected person is a woman, and 0 if the uninfected person is a man; $HSV2_{infected}$ is a binary factor which is 1 if the HIV infected person has HSV-2, and 0 otherwise; $HSV2_{uninfected}$ is a binary factor which is 1 if the HIV uninfected person has HSV-2, and 0 otherwise.

#### 3.2. Overview of StepSyn 1.0 parameters that were fitted to calibration targets

Table S5 gives an overview of the StepSyn 1.0 parameters that were fitted to the calibration targets in Table S3, together with the initial range ([minimum, maximum]) used for fitting.

**Table S5.** Overview of the StepSyn 1.0 parameters that were fitted to the calibration targets in Table S3. The last column indicates in which models these parameters were used (see main text for abbreviations).

| parameter definition | parameter name | initial range | models |
|---|---|---|---|
| HIV baseline probability male to female (m2f) (annotated by baseline in the figures) | hiv.tr.prob.m2f.baseline | [0.001, 0.02] | all |
| HIV transmission probability – influence of gender multiplier for female to male transmission (f2m) (annotated by fm_ratio in the figures) | hiv.tr.prob.baseline.f.m.ratio | [0.25, 0.5] | all |
| HIV transmission probability - influence of the HIV-infected person (index) being infected with HSV-2 (simplified HSV-2 options) (annotated by hsv2_index in the figures) | hiv.tr.prob.baseline.mult.hsv2.chronic.index | [1, 4] | St_Ba<br>St_IO |
| HIV transmission probability - influence of the HIV-uninfected person (exposed) being infected with HSV-2 (simplified HSV-2 options) (annotated by hsv2_exposed in the figures) | hiv.tr.prob.baseline.mult.hsv2.chronic.exposed | [1, 4] | St_Ba<br>St_IO |
| HIV transmission probability - influence of only the HIV-infected person (index) having an ulcerative recurrence of HSV-2 at the time (full HSV-2 options) | hiv.tr.prob.m2f.maleGUD<br>hiv.tr.prob.f2m.femaleGUD<br>These 2 parameters are assumed to have equal values. | [0.04, 0.1] | St_RG,<br>St_IO_RG |
| HIV transmission probability - influence of only the HIV-uninfected person (exposed) having an ulcerative recurrence of HSV-2 at the time (full HSV-2 options) | hiv.tr.prob.m2f.femaleGUD<br>hiv.tr.prob.f2m.maleGUD<br>These 2 parameters are assumed to have equal values. | [0.07, 0.1] | St_RG,<br>St_IO_RG |
| HIV transmission probability – influence of both the HIV-infected and the HIV-uninfected person having an ulcerative recurrence of HSV-2 at the time (full HSV-2 options) | hiv.tr.prob.m2f.bothGUD<br>hiv.tr.prob.f2m.bothGUD<br>These 2 parameters are assumed to have equal values. | [0.43, 0.5] | St_RG,<br>St_IO_RG |

##### 3.3. Parameter fitting methodology

The parameters for the Simpact Cyan 1.0 models (Table S4) and the StepSyn 1.0 models (Table S5) were fitted to the calibration targets in Table S3 by applying an iterative active learning approach [8] using the procedure described in [9] and minimizing the sum of squared relative errors [10] to determine model performance. The remaining model parameters were drawn from the literature (see paragraph 4 of this Supplementary Material).

###### *3.3.1. Latin hypercube sampling*

For each of the HIV transmission parameters, values were drawn from a uniform distribution, using the ranges from Table S4 and Table S5. These uniform distributions were used as initial (prior) probability distributions for the parameters in the parameter fitting procedure. We applied Latin Hypercube Sampling (LHS) [11] to select 10,000 parameter sets.

###### *3.3.2. Goodness-of-fit (GOF) statistic*

To find the values of the HIV transmission parameters most supported by the data in Table 3, we calculate the sum of squared relative errors [10] for each of the 10,000 parameter combinations.

###### *3.3.3. Statistical evaluation of the model parameters*

For the statistical evaluation of the model parameters, we focus on the subset of parameter combinations corresponding to the top 1% of the lowest values for the GOF statistic. In brief, the parameter fitting procedure consists of the following steps. First, a parameter wise comparison between the density of the initial probability distribution of the parameters and the density of the probability distribution of the subset of parameters corresponding with the top 1% solutions is conducted to determine which parameters are highly influenced by the data. Second, classification trees and generalized additive models are applied to determine which patterns of parameter vectors characterize the subspace of the top 1% solutions. Finally, we apply the Maximal Information Coefficient (MIC) [12] to determine associations between the parameters in the subspace of the top 1%.

Based on the results of the analyses above, we can narrow the solution space and repeat LHS and the steps described above several times.

A more detailed description of each step in the parameter fitting methodology, together with examples for Simpact Cyan 1.0 and StepSyn 1.0 is given below.

The R-scripts that have been used for fitting HIV transmission parameters are available from GitHub:

https://github.com/dmhendrickx/Scripts_comparison_Simpact_StepSyn

###### 3.3.3.1. Univariate explorative analysis – top 1% parameter combinations

To determine which parameters are highly influenced by the data, we compared the (prior) density of the initial uniform distribution (for the range in Tables S4 and S5) with the (posterior) density of the distribution of the top 1% parameter combinations for each parameter separately. A more peaked density for the top 1% parameter combinations indicates a parameter that is more influenced by the data.

*Example 1: Simpact Cyan 1.0 basic model (Si_Ba)*

If we apply univariate explorative analysis to the 10,000 LHS-generated combinations of parameters for the Si_Ba model, using the ranges in Table S4, we obtain the smoothed density plots shown in Figure S2. We observe that the baseline parameter (hiv_a) is the most influenced by the data. The remaining parameters are equally influenced by the data.

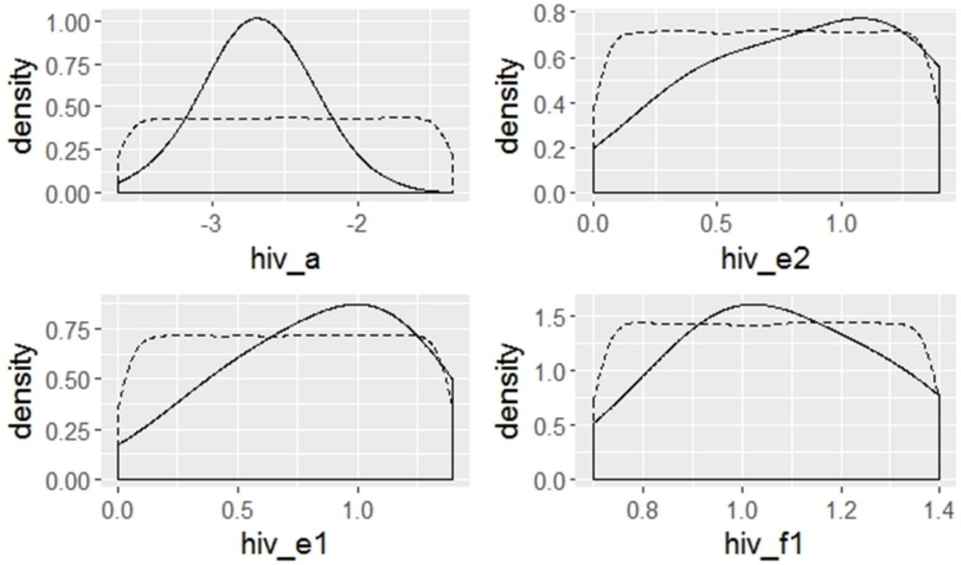


**Figure S2.** Simpact Cyan 1.0 basic model (Si_Ba). Overlaid smoothed density plots for the density of the initial uniform distribution of the parameters (dashed line) and density of the distribution of the top 1% parameter combinations (solid line). Parameters: hiv_a: HIV baseline transmission hazard a; hiv_f1: HIV transmission hazard – influence of gender; hiv_e1: HIV transmission hazard - influence of the HIV-infected person (index) being infected with HSV-2; hiv_e2: HIV transmission hazard - influence of the HIV-uninfected person (exposed) being infected with HSV-2.

A smoothed density plot is a smoothed histogram, e.g. the dashed line in the upper left panel of Figure S2 is a smoothed version of the histogram presented in Figure S3.

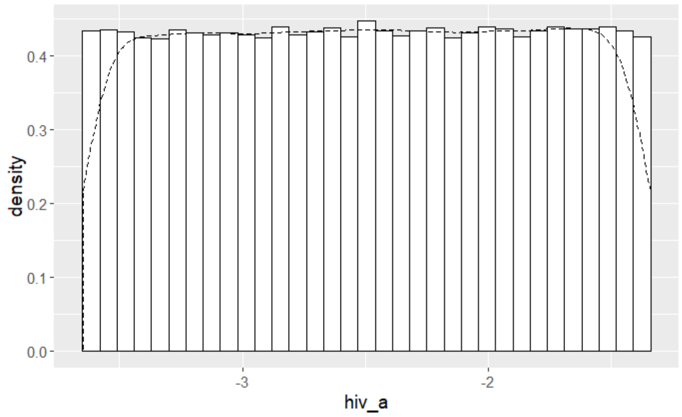


**Figure S3.** Histogram of the initial distribution of parameter hiv_a and smoothed density (dashed line).

*Example 2: StepSyn 1.0 basic model (St_Ba)*

If we apply univariate explorative analysis to the 10,000 LHS-generated combinations of parameters for the St_Ba model, using the ranges in Table S5, we obtain the smoothed density plots shown in Figure S5. We observe that the baseline parameter (baseline) is the most influenced by the data. The remaining parameters are equally influenced by the data.

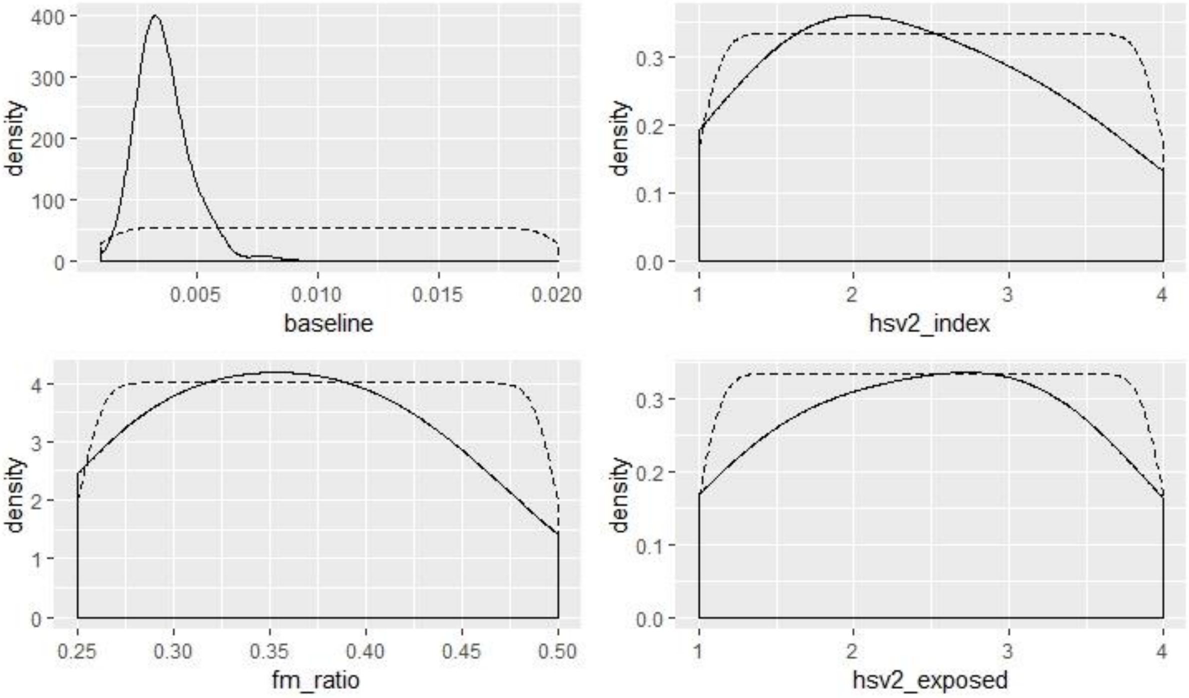


**Figure S4.** StepSyn 1.0 basic model (St_Ba). Overlaid smoothed density plots for the density of the initial uniform distribution of the parameters (dashed line) and density of the distribution of the top 1% parameter combinations (solid line). Parameters: baseline: HIV baseline probability; fm_ratio: HIV transmission probability – influence of gender; hsv2_index: HIV transmission probability - influence of the HIV-infected person (index) being infected with HSV-2 (simplified HSV-2 options); hsv2_exposed: HIV transmission probability - influence of the HIV-uninfected person (exposed) being infected with HSV-2 (simplified HSV-2 options).

3.3.3.2. Activity region finder - top 1% parameter combinations

To determine which patterns of parameter vectors characterize the subspace of top 1% solutions, we applied Activity Region Finder (ARF)[13], a recursive partitioning classification tree algorithm. We consider each of the 10,000 parameter vectors generated by LHS as input, and the binary variable that equals 1 for a parameter combination in the top 1% and 0 otherwise as output.

*Example 1: Simpact Cyan 1.0 basic model (Si_Ba)*

If we applied ARF to the 10,000 LHS-generated combinations of parameters for the Si_Ba model using the ranges in Table S4, we observed that the baseline parameter (hiv_a) is responsible for the first split of the tree, followed by splits using the influence of HIV-uninfected person (exposed) being infected with HSV-2 (hiv_e2) (left and middle branch of the tree) and the influence of gender (hiv_f1) (right branch of the tree). In total 7 regions were classified as significant high activity regions (= regions associated with a low sum of squared relative errors), see Table S6.

**Table S6.** Simpact Cyan 1.0 basic model (Si_Ba). Parameter regions classified as high activity regions by ARF. Parameters: baseline: HIV baseline probability; fm_ratio: HIV transmission probability – influence of gender; hsv2_index: HIV transmission probability - influence of the HIV-infected person (index) being infected with HSV-2 (simplified HSV-2 options); hsv2_exposed: HIV transmission probability - influence of the HIV-uninfected person (exposed) being infected with HSV-2 (simplified HSV-2 options). L = left branch; M = middle branch; R = right branch.

| Region | Parameter ranges |
|---|---|
| M | hiv_a in [-3.0503,-2.2737] |
| MRM | hiv_a in [-3.0503,-2.2737]; hiv_e2 in [1.2851,1.3999]; hiv_e1 in [0.4607,0.6832] |
| MLRM | hiv_a in [-3.0503,-2.2737]; hiv_e2 in [5e-04,1.2638]; hiv_e1 in [1.256,1.3997]; hiv_f1 in [1.0051,1.1011] |
| MLLLM | hiv_a in [-2.2953,-2.2737]; hiv_e2 in [5e-04,1.2638]; hiv_e1 in [0,0.8981] |
| MLLRM | hiv_a in [-3.0503,-2.2737]; hiv_e2 in [5e-04,1.2638]; hiv_e1 in [1.0294,1.0465] |
| MLLLLM | hiv_a in [-3.0503,-2.2954]; hiv_e2 in [5e-04,1.2638]; hiv_e1 in [0.7001,0.8669] |
| MLLRRM | hiv_a in [-3.0503,-2.2737]; hiv_e2 in [0.4281,0.596]; hiv_e1 in [1.0472,1.2286] |

*Example 2: StepSyn 1.0 basic model (St_Ba)*

If we applied ARF to the 10,000 LHS-generated combinations of parameters for the St_Ba model using the ranges in Table S5, we observed that the baseline parameter (baseline) is responsible for the first two splits of the tree, followed by splits using the influence of HIV-uninfected person (exposed) being infected with HSV-2 (hsv2_exposed) (middle left and middle right branch of the tree) and the influence of gender (fm_ratio) (right middle branch of the tree). In total 7 regions were classified as significant high activity regions (= regions associated with a low sum of squared relative errors), see Table S7.

**Table S7.** StepSyn 1.0 basic model (St_Ba). Parameter regions classified as high activity regions by ARF. Parameters: baseline: HIV baseline probability; fm_ratio: HIV transmission probability – influence of gender; hsv2_index: HIV transmission probability - influence of the HIV-infected person (index) being infected with HSV-2 (simplified HSV-2 options); hsv2_exposed: HIV transmission probability - influence of the HIV-uninfected person (exposed) being infected with HSV-2 (simplified HSV-2 options). L = left branch; M = middle branch; R = right branch.

| Region | Parameter ranges |
|---|---|
| M | baseline in [0.002297,0.004016] |
| RM | baseline in [0.004072,0.005759] |
| MRM | baseline in [0.003565,0.004016] |
| RMM | baseline in [0.004072,0.005759]; fm_ratio in [0.26116,0.26569] |
| MLMM | baseline in [0.002851,0.0030826]; hsv2_exposed in [2.941,3.939] |
| MLLLM | baseline in [0.0023,0.003186]; hsv2_exposed in [1.0087,2.9393]; fm_ratio in [0.3742,0.3945] |
| RMRMM | baseline in [0.004072,0.005759]; fm_ratio in [0.266,0.4998]; hsv2_index in [1.317,1.8496]; hsv2_exposed in [1.4206,1.795] |

3.3.3.3. Generalized additive models - top 1% parameter combinations

As a second method to determine which patterns of parameter vectors characterize the subspace of top 1% solutions, we applied generalized additive models (GAM)[14] considering the same input and output variables as used with ARF.

For selecting the tuning parameter of the GAM, we consider both the Akaike Information Criterion (AIC)[15] and the Bayesian Information Criterion (BIC)[16].

*Example 1: Simpact Cyan 1.0 basic model (Si_Ba)*

If we applied GAM to the 10,000 LHS-generated combinations of parameters for the Si_Ba model using the ranges in Table S4, the models with tuning parameter 1 and 10 had the lowest AIC and BIC respectively, see Figure S5.

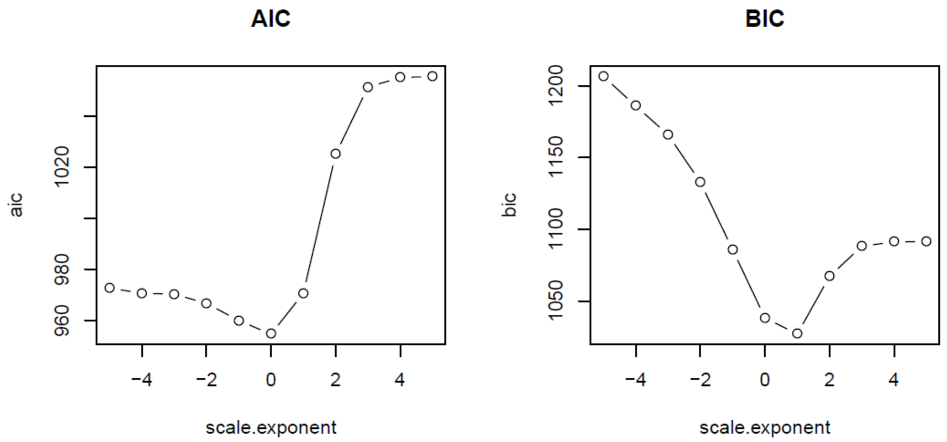


**Figure S5.** Simpact Cyan 1.0 basic model (Si_Ba). Selection of the tuning parameter for the GAM based on AIC and BIC. Scale.exponent = 10log(tuning parameter).

The figures for both GAMs (Figures S6 and S7) show that intermediate values of the baseline parameter (hiv_a) are associated with a higher probability of obtaining a low relative sum of squared errors than low and high values. For the influence of the HIV-infected person (index) being infected with HSV-2 (hiv_e1) and the influence of the HIV-uninfected person (exposed) being infected with HSV-2 (hiv_e2), the probability of obtaining a low relative sum of squared errors increases with the parameter value.

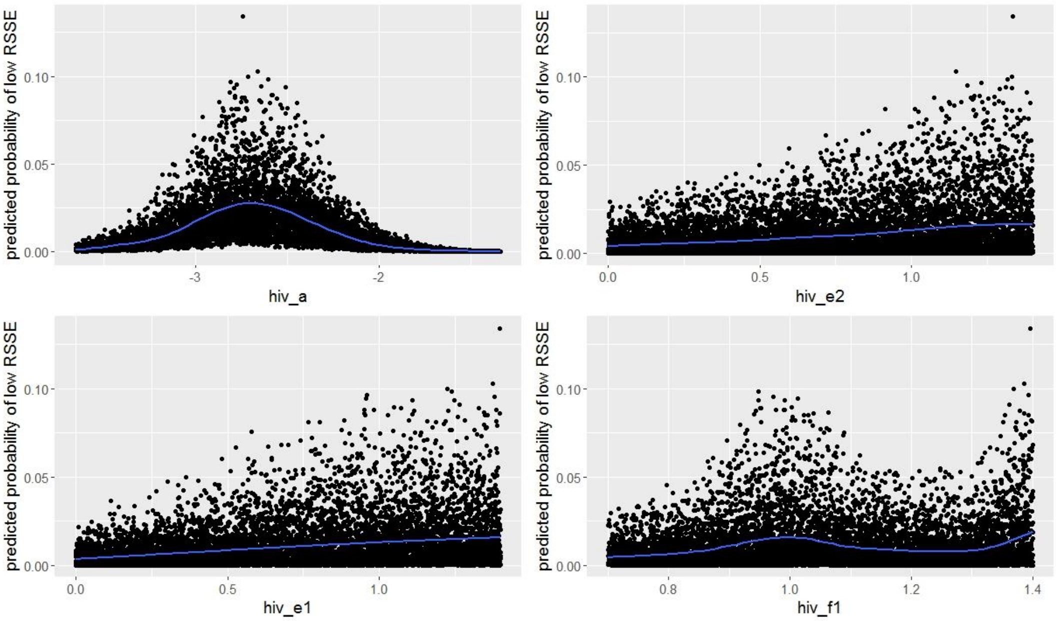


**Figure S6.** Simpact Cyan 1.0 basic model (Si_Ba). Results of the generalized additive model, with the tuning parameter selected by AIC. Predicted probabilities of a low relative sum of squared errors (RSSE) for 10,000 LHS-generated combinations of Simpact Cyan 1.0 parameters using the ranges in Table S4. Parameters: hiv_a: HIV baseline transmission hazard a; hiv_f1: HIV transmission hazard – influence of gender; hiv_e1: HIV transmission hazard - influence of the HIV-infected person (index) being infected with HSV-2; hiv_e2: HIV transmission hazard - influence of the HIV-uninfected person (exposed) being infected with HSV-2.

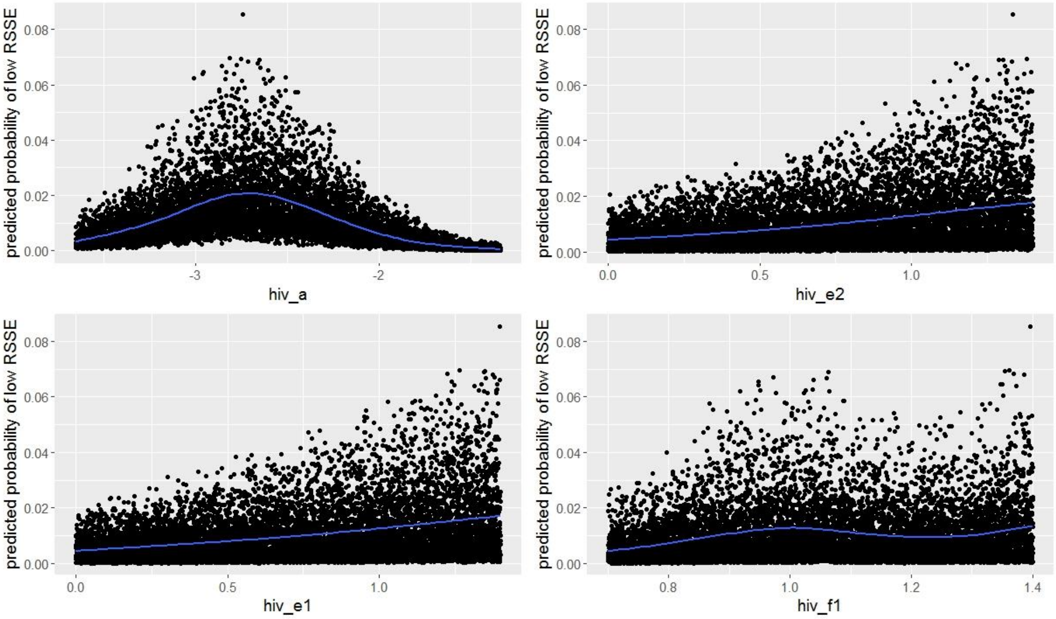


**Figure S7.** Simpact Cyan 1.0 basic model (Si_Ba). Results of the generalized additive model, with tuning parameter selected by BIC. Predicted probabilities of low relative sum of squared errors (RSSE) for 10,000 LHS-generated combinations of Simpact Cyan 1.0 parameters using the ranges in Table S4. Parameters: hiv_a: HIV baseline transmission hazard a; hiv_f1: HIV transmission hazard – influence of gender; hiv_e1: HIV transmission hazard - influence of the HIV-infected person (index) being infected with HSV-2; hiv_e2: HIV transmission hazard - influence of the HIV-uninfected person (exposed) being infected with HSV-2.

*Example 2: StepSyn 1.0 basic model (St_Ba)*

If we applied GAM to the 10,000 LHS-generated combinations of parameters for the St_Ba model using the ranges in Table S5, the models with tuning parameter 1 and 10 had the lowest AIC and BIC respectively, see Figure S8.

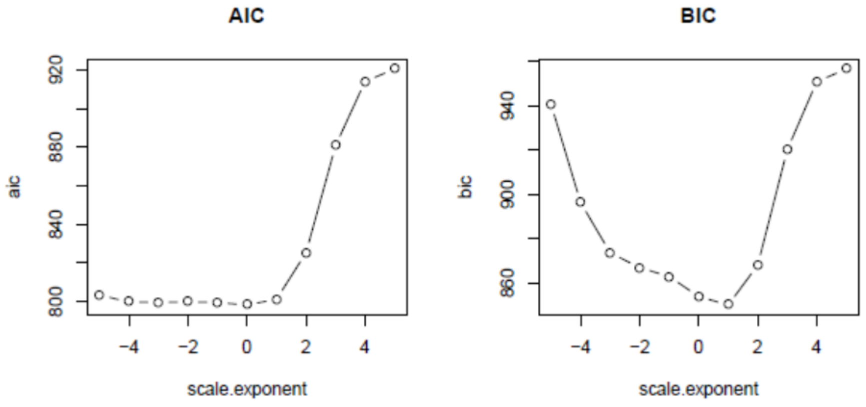


**Figure S8.** StepSyn 1.0 basic model (St_Ba). Selection of the tuning parameter for the GAM based on AIC and BIC. Scale.exponent = 10log(tuning parameter).

The figures for both GAMs (Figures S9 and S10) show that low values of the baseline parameter are associated with a higher probability of obtaining a low relative sum of squared errors than intermediate and high values. For the influence of gender (fm_ratio) and influence of the HIV-infected person (index) being infected with HSV-2 (hsv2_index), the probability of obtaining a low relative sum of squared errors increases with the parameter value.

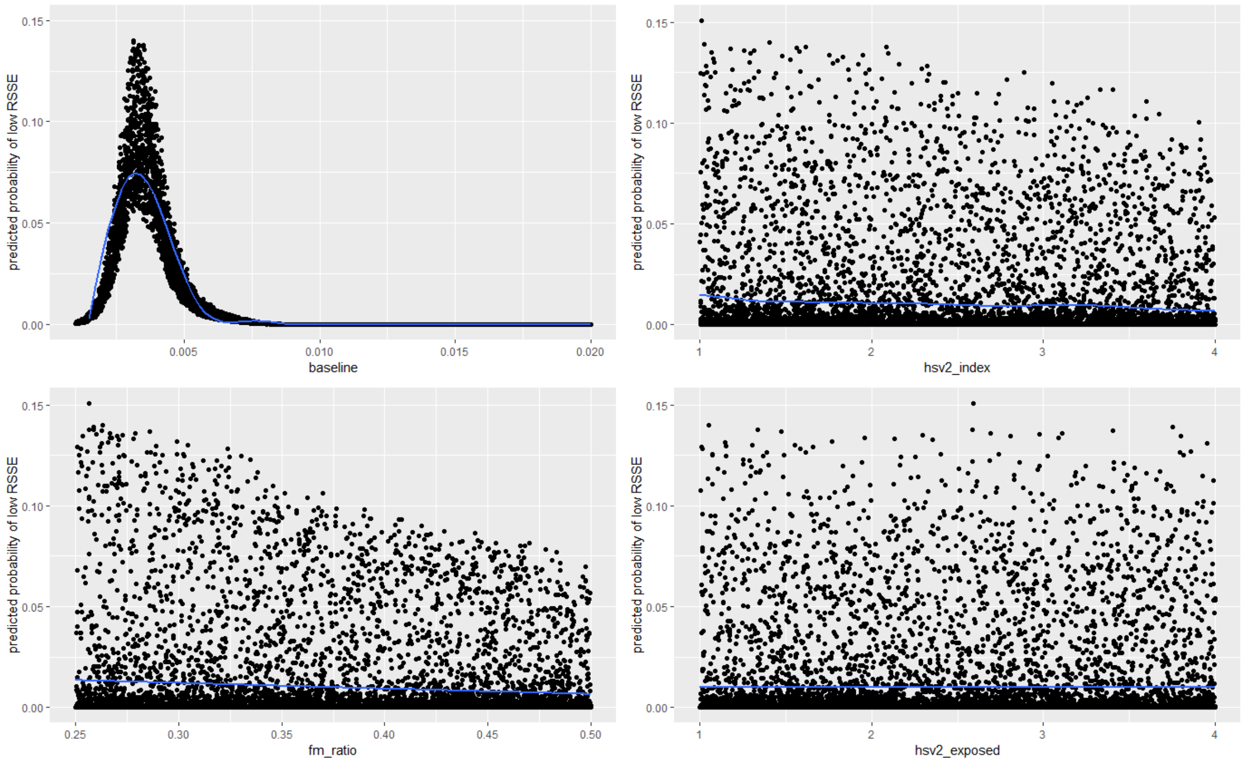


**Figure S9.** StepSyn 1.0 basic model (St_Ba). Results of the generalized additive model, with tuning parameter selected by AIC. Predicted probabilities of low relative sum of squared errors (RSSE) for 10,000 LHS-generated combinations of StepSyn 1.0 parameters using the ranges in Table S5. Parameters: baseline: HIV baseline probability; fm_ratio: HIV transmission probability – influence of gender; hsv2_index: HIV transmission probability - influence of the HIV-infected person (index) being infected with HSV-2 (simplified HSV-2 options); hsv2_exposed: HIV transmission probability - influence of the HIV-uninfected person (exposed) being infected with HSV-2 (simplified HSV-2 options).

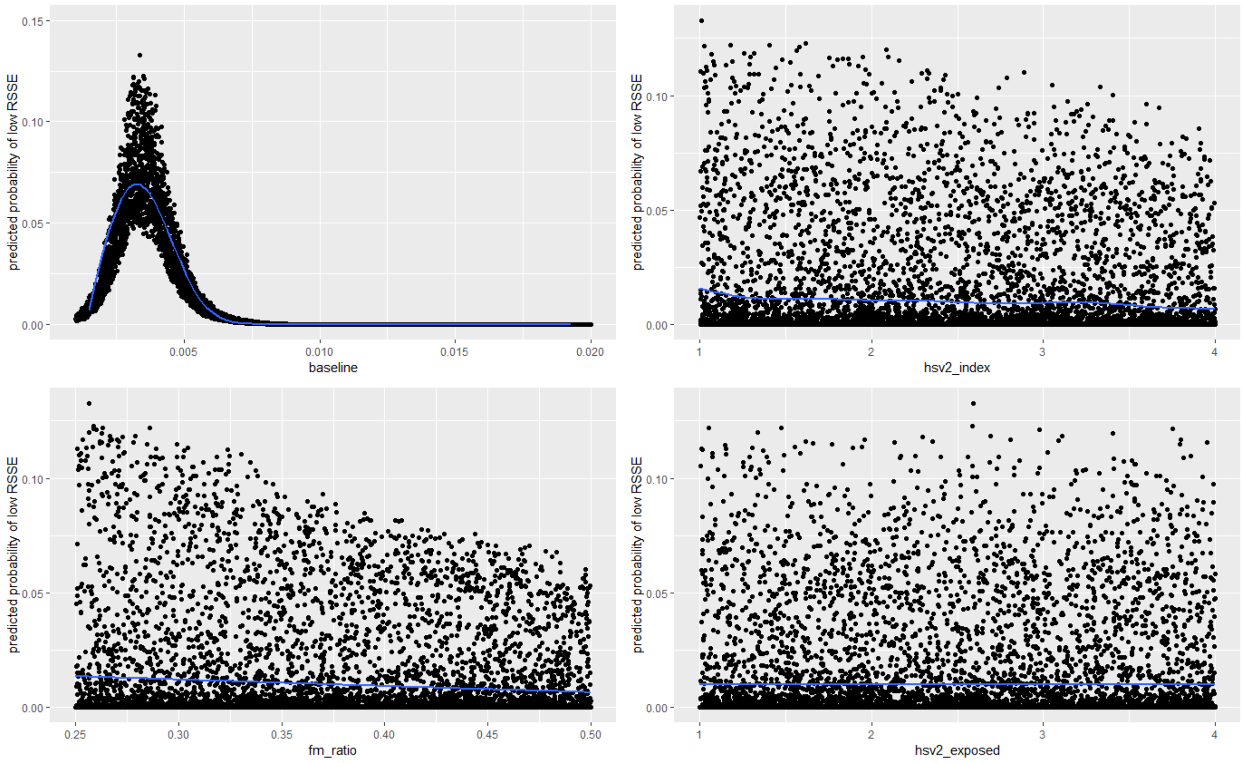


**Figure S10.** StepSyn 1.0 basic model (St_Ba). Results of the generalized additive model, with tuning parameter selected by BIC. Predicted probabilities of low relative sum of squared errors (RSSE) for 10,000 LHS-generated combinations of StepSyn 1.0 parameters using the ranges in Table S5. Parameters: baseline: HIV baseline probability; fm_ratio: HIV transmission probability – influence of gender; hsv2_index: HIV transmission probability - influence of the HIV-infected person (index) being infected with HSV-2 (simplified HSV-2 options); hsv2_exposed: HIV transmission probability - influence of the HIV-uninfected person (exposed) being infected with HSV-2 (simplified HSV-2 options).

3.3.3.4. Maximal Information Coefficient - top 1% parameter combinations

To determine associations between the parameters within the subset of the top 1% parameter combinations, we applied the Maximal Information Coefficient (MIC)[12].

*Example 1: Simpact Cyan 1.0 basic model (Si_Ba)*

Table S8 shows the results for the top 1% of 10,000 LHS-generated combinations of parameters for the Si_Ba model using the ranges in Table S4. We observed the highest value of the MIC for the association between the baseline parameter (hiv_a) and the parameter for the influence of the HIV-uninfected person being infected with HSV-2 (hiv_e2). The second highest value of the MIC was observed for the association between the baseline parameter (hiv_a) and the parameter for the influence of the HIV-infected person being infected with HSV-2 (hiv_e1). Figure S11 shows that both associations are negative. All other associations had a MIC smaller than 0.35.

**Table S8.** Simpact Cyan 1.0 basic model (Si_Ba). MIC for the top 1% of 10,000 LHS-generated combinations of Simpact Cyan 1.0 parameters using the ranges in Table S4. Parameters: hiv_a: HIV baseline transmission hazard a; hiv_f1: HIV transmission hazard – influence of gender; hiv_e1: HIV transmission hazard - influence of the HIV-infected person (index) being infected with HSV-2; hiv_e2: HIV transmission hazard - influence of the HIV-uninfected person (exposed) being infected with HSV-2.

| parameter 1 | parameter 2 | MIC |
|---|---|---|
| hiv_a | hiv_e1 | 0.390463 |
| hiv_a | hiv_e2 | 0.486962 |
| hiv_a | hiv_f1 | 0.3227297 |
| hiv_e1 | hiv_e2 | 0.2094048 |
| hiv_e1 | hiv_f1 | 0.287366 |
| hiv_e2 | hiv_f1 | 0.2704736 |

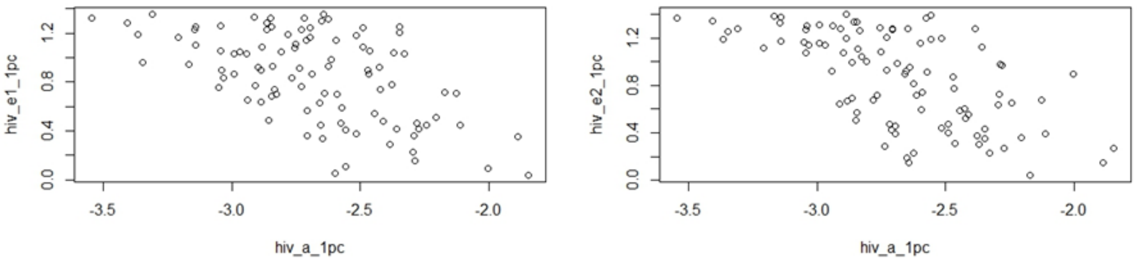


**Figure S11.** Simpact Cyan 1.0 basic model (Si_Ba). Left: hiv_e1 vs. hiv_a for the top 1% parameter combinations. Right: hiv_e2 vs. hiv_a. 1pc: top 1% solutions. Parameters: hiv_a: HIV baseline transmission hazard a; hiv_e1: HIV transmission hazard - influence of the HIV-infected person (index) being infected with HSV-2; hiv_e2: HIV transmission hazard - influence of the HIV-uninfected person (exposed) being infected with HSV-2.

*Example 2: StepSyn 1.0 basic model (St_Ba)*

Table S9 shows the results for the top 1% of 10,000 LHS-generated combinations of parameters for the St_Ba model using the ranges in Table S5. The results show that all associations had a MIC smaller than 0.3.

**Table S9.** StepSyn 1.0 basic model (St_Ba). MIC for the top 1% of 10,000 LHS-generated combinations of StepSyn 1.0 parameters using the ranges in Table S5. Parameters: baseline: HIV baseline probability; fm_ratio: HIV transmission probability – influence of gender; hsv2_index: HIV transmission probability - influence of the HIV-infected person (index) being infected with HSV-2 (simplified HSV-2 options); hsv2_exposed: HIV transmission probability - influence of the HIV-uninfected person (exposed) being infected with HSV-2 (simplified HSV-2 options).

| parameter 1 | parameter 2 | MIC |
|---|---|---|
| baseline | fmratio | 0 |
| baseline | hsv2_index | 0 |
| baseline | hsv2_exposed | 0 |
| fmratio | hsv2_index | 0.2062702 |
| fmratio | hsv2_exposed | 0.1976175 |
| hsv2_index | hsv2_exposed | 0.2626614 |

##### *3.3.4. Active learning*

From the results of the statistical evaluation of the parameters described in 3.3.3, we determine new ranges for the parameters and repeat the steps described in 3.3.1, 3.3.2 and 3.3.3. We repeat this several times until the solution cannot be improved anymore.

This approach is called iterative active learning [8]. In most cases, 3-4 iterations are sufficient to determine the final solution for the parameters.

In our study, the new ranges for the parameters were chosen so that they include

- the parameter combination with the lowest RSSE

- the significant high activity regions determined by ARF

- results from GAM (only in case they define smaller intervals for some parameters than ARF)

*Example 1: Simpact Cyan 1.0 basic model (Si_Ba)*

Table S10 shows how we determined the parameter ranges for the Si_Ba model for the second iteration in the active learning approach.

**Table S10.** Simpact Cyan 1.0 basic model (Si_Ba). Selecting of a new range of Simpact Cyan 1.0 parameters from the results in paragraph 3.3.3. Parameters: hiv_a: HIV baseline transmission hazard a; hiv_f1: HIV transmission hazard – influence of gender; hiv_e1: HIV transmission hazard - influence of the HIV-infected person (index) being infected with HSV-2; hiv_e2: HIV transmission hazard - influence of the HIV-uninfected person (exposed) being infected with HSV-2. RSSE = relative sum of squared errors; ARF = activity region finder.

| Parameter | Solution with lowest RSSE | ARF | New range |
|---|---|---|---|
| hiv_a | -2.864607 | [-3.06, -2.27] | [-3.06, -2.27] |
| hiv_f1 | 1.382483 | [1, 1.11] | [1, 1.4] |
| hiv_e1 | 1.286708 | [0, 1.4] | [0, 1.4] |
| hiv_e2 | 0.994065 | [0, 1.4] | [0, 1.4] |

*Example 2: StepSyn 1.0 basic model (St_Ba)*

Table S11 shows how we determined the parameter ranges for the St_Ba model for the second iteration in the active learning approach.

**Table S11.** StepSyn 1.0 basic model (St_Ba). Selecting of a new range of StepSyn 1.0 parameters from the results in paragraph 3.3.3. Parameters: baseline: HIV baseline probability; fm_ratio: HIV transmission probability – influence of gender; hsv2_index: HIV transmission probability - influence of the HIV-infected person (index) being infected with HSV-2 (simplified HSV-2 options); hsv2_exposed: HIV transmission probability - influence of the HIV-uninfected person (exposed) being infected with HSV-2 (simplified HSV-2 options). RSSE = relative sum of squared errors; ARF = activity region finder.

| Parameter | Solution with lowest RSSE | ARF | New range |
|---|---|---|---|
| baseline | 0.004534496 | [0.002, 0.006] | [0.002, 0.006] |
| fm_ratio | 0.383881350 | [0.26, 0.50] | [0.26, 0.50] |
| hsv2_index | 1.778933848 | [1.31, 1.85] | [1.31, 1.85] |
| hsv2_exposed | 1.795042948 | [1, 4] | [1, 4] |

*Determination of final parameters*

Table S12 shows that both for the Simpact Cyan 1.0 basic model (Si_Ba) and the StepSyn 1.0 basic model (St_Ba), we could determine the final parameters (see Supplementary Material, paragraph 4) after three iterations of active learning (the parameters corresponding with the lowest RSSE in iteration 2).

**Table S12.** Lowest RSSE for Simpact Cyan 1.0 basic model (Si_Ba) and StepSyn 1.0 basic model (St_Ba) for three iterations of active learning.

| Iteration | Simpact Cyan 1.0 - Lowest RSSE | StepSyn – Lowest RSSE |
|---|---|---|
| 1 | 1.279048 | 1.291408 |
| 2 | 1.184201 (best) | 0.9940927 (best) |
| 3 | 1.21711 | 1.34732 |

### 4. Estimated and other key model parameters

#### 4.1. Simpact Cyan 1.0 basic model (Si_Ba)

**Table S13.** Estimated and other key model parameters for the Simpact Cyan 1.0 basic model (Si_Ba).

| parameter definition | parameter name | parameter value | source / reference |
|---|---|---|---|
| HIV baseline transmission hazard a | hivtransmission.param.a | -3.01 | estimated in this study |
| HIV transmission hazard - influence of the HIV-infected person (index) being infected with HSV-2 | hivtransmission.param.e1 | 0.89 | estimated in this study |
| HIV transmission hazard - influence of the HIV-uninfected person (exposed) being infected with HSV-2 | hivtransmission.param.e2 | 1.37 | estimated in this study |
| HIV transmission hazard – influence of gender | hivtransmission.param.f1 | 1.31 | estimated in this study |
| HSV-2 baseline transmission hazard* | person.hsv2.a.dist.fixed.value | -2.55 | estimated in this study, so that the seroprevalence of HSV-2 first increases and afterwards stabilizes at approximately 50% for females [17] |

* Transmission probability from the literature converted to a transmission hazard using the procedure described in Supplementary Material, paragraph 4.11.

#### 4.2. Simpact Cyan 1.0 model with inflow and outflow (Si_IO).

**Table S14.** Estimated and other key model parameters for the Simpact Cyan 1.0 model with inflow and outflow (Si_IO).

| parameter definition | parameter name | parameter value | source / reference |
|---|---|---|---|
| HIV baseline transmission hazard a | hivtransmission.param.a | -2.29 | estimated in this study |
| HIV transmission hazard - influence of the HIV-infected person (index) being infected with HSV-2 | hivtransmission.param.e1 | 1.03 | estimated in this study |
| HIV transmission hazard - influence of the HIV-uninfected person (exposed) being infected with HSV-2 | hivtransmission.param.e2 | 1.22 | estimated in this study |
| HIV transmission hazard – influence of gender | hivtransmission.param.f1 | 0.97 | estimated in this study |
| HSV-2 baseline transmission hazard* | person.hsv2.a.dist.fixed.value | -1.5 | estimated in this study, so that the seroprevalence of HSV-2 first increases and afterwards stabilizes at approximately 50% for females [17] |
| non-AIDS mortality – shape parameter for Weibull distribution | mortality.normal.weibull.shape | 1.8 | estimated from population pyramid Yaoundé [18] |
| non-AIDS mortality – scale parameter for Weibull distribution | mortality.normal.weibull.scale | 52 | estimated from population pyramid Yaoundé [18] |
| non-AIDS mortality – influence of gender - for a woman, half this value is added to the scale parameter of the Weibull distribution. For a man, the same amount is subtracted. | mortality.normal.weibull.genderdiff | 2 | estimated from population pyramid Yaoundé [18] |

* Transmission probability from the literature converted to a transmission hazard using the procedure described in Supplementary Material, paragraph 4.11.

#### 4.3. Simpact Cyan 1.0 model implementing viral load (Si_VL).

**Table S15.** Estimated and other key model parameters for the Simpact Cyan 1.0 model implementing viral load (Si_VL).

| parameter definition | parameter name | parameter value | source / reference |
|---|---|---|---|
| HIV baseline transmission hazard a | hivtransmission.param.a | -1.65 | estimated in this study |
| HIV transmission hazard – parameter b in the formula $hazard = \exp\,(a + bV^{-c})$ [7] where a = baseline transmission hazard (see above) V = current viral load | hivtransmission.param.b | - 12.49 | estimated in this study |
| HIV transmission hazard – parameter c in the formula $hazard = \exp\,(a + bV^{-c})$ [7] where a = baseline transmission hazard (see above) V = current viral load | hivtransmission.param.c | 0.20 | estimated in this study |
| HIV transmission hazard - influence of the HIV-infected person (index) being infected with HSV-2 | hivtransmission.param.e1 | 1.31 | estimated in this study |
| HIV transmission hazard - influence of the HIV-uninfected person (exposed) being infected with HSV-2 | hivtransmission.param.e2 | 1.32 | estimated in this study |
| HIV transmission hazard – influence of gender | hivtransmission.param.f1 | 1.20 | estimated in this study |
| HSV-2 baseline transmission hazard* | person.hsv2.a.dist.fixed.value | -2.55 | estimated in this study, so that the seroprevalence of HSV-2 first increases and afterwards stabilizes at approximately 50% for females [17] |

* Transmission probability from the literature converted to a transmission hazard using the procedure described in Supplementary Material, paragraph 4.11.

#### 4.4. Simpact Cyan 1.0 model implementing inflow, outflow and viral load (Si_IO_VL).

**Table S16.** Estimated and other key model parameters for the Simpact Cyan 1.0 model implementing inflow, outflow and viral load (Si_IO_VL).

| parameter definition | parameter name | parameter value | source / reference |
|---|---|---|---|
| HIV baseline transmission hazard a | hivtransmission.param.a | -1.59 | estimated in this study |
| HIV transmission hazard – parameter b in the formula $hazard = \exp(a + bV^{-c})$ [7] where a = baseline transmission hazard (see above) V = current viral load | hivtransmission.param.b | -10.66 | estimated in this study |
| HIV transmission hazard – parameter c in the formula $hazard = \exp(a + bV^{-c})$ [7] where a = baseline transmission hazard (see above) V = current viral load | hivtransmission.param.c | 0.29 | estimated in this study |
| HIV transmission hazard - influence of the HIV-infected person (index) being infected with HSV-2 | hivtransmission.param.e1 | 1.11 | estimated in this study |
| HIV transmission hazard - influence of the HIV-uninfected person (exposed) being infected with HSV-2 | hivtransmission.param.e2 | 0.98 | estimated in this study |
| HIV transmission hazard – influence of gender | hivtransmission.param.f1 | 0.89 | estimated in this study |
| HSV-2 baseline transmission hazard* | person.hsv2.a.dist.fixed.value | -1.5 | estimated in this study, so that the seroprevalence of HSV-2 first increases and afterwards stabilizes at approximately 50% for females [17] |
| non-AIDS mortality – shape parameter for Weibull distribution | mortality.normal.weibull.shape | 1.8 | estimated from population pyramid Yaoundé [18] |
| non-AIDS mortality – scale parameter for Weibull distribution | mortality.normal.weibull.scale | 52 | estimated from population pyramid Yaoundé [18] |
| non-AIDS mortality – influence of gender - for a woman, half this value is added to the scale parameter of the Weibull distribution. For a man, the same amount is subtracted. | mortality.normal.weibull.genderdiff | 2 | estimated from population pyramid Yaoundé [18] |

* Transmission probability from the literature converted to a transmission hazard using the procedure described in Supplementary Material, paragraph 4.11.

#### 4.5. StepSyn 1.0 basic model (St_Ba)

**Table S17.** Estimated and other key model parameters for the StepSyn 1.0 basic model (St_Ba).

| parameter definition | parameter name | parameter value | source / reference |
|---|---|---|---|
| HIV baseline probability | hiv.tr.prob.m2f.baseline | 0.0045 | estimated in this study |
| HIV transmission probability – influence of gender | hiv.tr.prob.baseline.f.m.ratio | 0.30 | estimated in this study |
| HIV transmission probability - influence of the HIV-infected person (index) being infected with HSV-2 (simplified HSV-2 options) | hiv.tr.prob.baseline.mult.hsv2.chronic.index | 1.37 | estimated in this study |
| HIV transmission probability - influence of the HIV-uninfected person (exposed) being infected with HSV-2 (simplified HSV-2 options) | hiv.tr.prob.baseline.mult.hsv2.chronic.exposed | 2.64 | estimated in this study |
| HSV-2 baseline transmission probability (simplified HSV-2 options) | hsv2.tr.prob.chronic.m2f | 0.003 | estimated in this study, so that the seroprevalence of HSV-2 first increases and afterwards stabilizes at approximately 50% for females [17] |

#### 4.6. StepSyn 1.0 model with inflow and outflow (St_IO)

**Table S18.** Estimated and other key model parameters for the StepSyn 1.0 model with inflow and outflow (St_IO).

| parameter definition | parameter name | parameter value | source / reference |
|---|---|---|---|
| HIV baseline probability | hiv.tr.prob.m2f.baseline | 0.0068 | estimated in this study |
| HIV transmission probability – influence of gender | hiv.tr.prob.baseline.f.m.ratio | 0.33 | estimated in this study |
| HIV transmission probability - influence of the HIV-infected person (index) being infected with HSV-2 (simplified HSV-2 options) | hiv.tr.prob.baseline.mult.hsv2.chronic.index | 1.79 | estimated in this study |
| HIV transmission probability - influence of the HIV-uninfected person (exposed) being infected with HSV-2 (simplified HSV-2 options) | hiv.tr.prob.baseline.mult.hsv2.chronic.exposed | 1.51 | estimated in this study |
| HSV-2 baseline transmission probability (simplified HSV-2 options) | hsv2.tr.prob.chronic.m2f | 0.01 | estimated in this study, so that the seroprevalence of HSV-2 first increases and afterwards stabilizes at approximately 50% for females [17] |
| birth rate | natality.rate | 0.036 | birth rate Cameroon: 3.6%* |
| (non-AIDS) mortality rate | mortality.rate | 0.01 | mortality rate Cameroon: 1%* |
| immigration rate | immigration.rate | 0.142 | popul growth in 1997: 6.8% [4]; assume emigr ~= 10%; gives immigr = 14.2% to obtain growth = 6.8% |
| emigration rate | emigration.rate | 0.1 | popul growth in 1997: 6.8% [4]; assume emigr ~= 10%; gives immigr = 14.2% to obtain growth = 6.8% |

*****https://www.indexmundi.com/cameroon/demographics_profile.html

#### 4.7. StepSyn 1.0 model with the full set of HSV-2 co-factor assumptions (St_RG).

**Table S19.** Estimated and other key model parameters for the StepSyn 1.0 model with the full set of HSV-2 co-factor assumptions (St_RG).

| parameter definition | parameter name | parameter value | source / reference |
|---|---|---|---|
| HIV baseline probability | hiv.tr.prob.m2f.baseline | 0.0025 | estimated in this study |
| HIV transmission probability – influence of gender | hiv.tr.prob.baseline.f.m.ratio | 0.44 | estimated in this study |
| HIV transmission probability - influence of only the HIV-infected person (index having an ulcerative recurrence of HSV-2 at the time (full HSV-2 options) | hiv.tr.prob.m2f.maleGUD hiv.tr.prob.f2m.femaleGUD These 2 parameters are assumed to have equal values. | 0.05 | estimated in this study |
| HIV transmission probability - influence of only the HIV-uninfected person (exposed) having an ulcerative recurrence of HSV-2 at the time (full HSV-2 options) | hiv.tr.prob.m2f.femaleGUD hiv.tr.prob.f2m.maleGUD These 2 parameters are assumed to have equal values. | 0.07 | estimated in this study |
| HIV transmission probability – influence of both the HIV-infected and the HIV-uninfected person having an ulcerative recurrence of HSV-2 at the time (full HSV-2 options) | hiv.tr.prob.m2f.bothGUD hiv.tr.prob.f2m.bothGUD These 2 parameters are assumed to have equal values. | 0.46 | estimated in this study |
| HSV-2 baseline transmission probability (full HSV-2 options) | hsv2.tr.prob.primary.m2f | 0.08 | estimated in this study, so that the seroprevalence of HSV-2 first increases and afterwards stabilizes at approximately 50% for females [17] |

#### 4.8. StepSyn 1.0 model with inflow, outflow and the full set of HSV-2 co-factor assumptions (St_IO_RG)

**Table S20.** Estimated and other key model parameters for the StepSyn 1.0 model with inflow, outflow and the full set of HSV-2 co-factor assumptions (St_IO_RG).

| parameter definition | parameter name | parameter value | source / reference |
|---|---|---|---|
| HIV baseline probability | hiv.tr.prob.m2f.baseline | 0.002 | estimated in this study |
| HIV transmission probability – influence of gender | hiv.tr.prob.baseline.f.m.ratio | 0.38 | estimated in this study |
| HIV transmission probability - influence of only the HIV-infected person (index) having an ulcerative recurrence of HSV-2 at the time (full HSV-2 options) | hiv.tr.prob.m2f.maleGUD<br>hiv.tr.prob.f2m.femaleGUD<br>These 2 parameters are assumed to have equal values. | 0.06 | estimated in this study |
| HIV transmission probability - influence of only the HIV-uninfected person (exposed) having an ulcerative recurrence of HSV-2 at the time (full HSV-2 options) | hiv.tr.prob.m2f.femaleGUD<br>hiv.tr.prob.f2m.maleGUD<br>These 2 parameters are assumed to have equal values. | 0.08 | estimated in this study |
| HIV transmission probability – influence of both the HIV-infected and the HIV-uninfected person having an ulcerative recurrence of HSV-2 at the time (full HSV-2 options) | hiv.tr.prob.m2f.bothGUD<br>hiv.tr.prob.f2m.bothGUD<br>These 2 parameters are assumed to have equal values. | 0.48 | estimated in this study |
| HSV-2 baseline transmission probability (full HSV-2 options) | hsv2.tr.prob.primary.m2f | 0.18 | estimated in this study, so that the seroprevalence of HSV-2 first increases and afterwards stabilizes at approximately 50% for females [17] |
| birth rate | natality.rate | 0.036 | birth rate Cameroon: 3.6%* |
| (non-AIDS) mortality rate | mortality.rate | 0.01 | mortality rate Cameroon: 1%* |
| immigration rate | immigration.rate | 0.142 | popul growth in 1997: 6.8% [4]; assume emigr ~= 10%; gives immigr = 14.2% to obtain growth = 6.8% |
| emigration rate | emigration.rate | 0.1 | popul growth in 1997: 6.8% [4]; assume emigr ~= 10%; gives immigr = 14.2% to obtain growth = 6.8% |

*****https://www.indexmundi.com/cameroon/demographics_profile.html

#### 4.9. Other parameters used in all Simpact Cyan 1.0 models

**Table S21.** Model parameters used in all Simpact Cyan 1.0 models in this study.

| parameter definition | parameter name | parameter value | source / reference |
|---|---|---|---|
| time frame of the simulation (years) | population.simtime | 35 | stabilization period of 10 years + 1980-2005 |
| duration acute stage | chronicstage.acutestagetime | 12/52 | 12 weeks [19] |
| time to AIDS: time before the AIDS-related death a person will advance to the AIDS stage of infection. | aidsstage.start | 1 | 1 year [20,21] |
| time of survival from AIDS | mortality.aids.survtime.k<br>mortality.aids.survtime.C | 0<br>2 | survival time = time to AIDS + 1 year [20,22] |
| HSV-2 transmission – influence of gender* | hsv2transmission.hazard.c | 0.694 | [23,24] |
| HIV seed in 1980 | hivseed.amount | for fitting: hivseed.amount equal to 0.5% of the population<br>100 simulations: seed drawn from a binomial distribution with probability = 0.005 | leads to an HIV prevalence of 0.6% for males and 1.2% for females in 1989 (values in Table S3) |
| HSV2 seed in 1980 | hsv2seed.fraction | 0.25 | [25] |

* Transmission probability from the literature converted to a transmission hazard using the procedure described in Supplementary Material, paragraph 4.11.

#### 4.10. Other parameters used in all StepSyn 1.0 models

**Table S22.** Model parameters used in all StepSyn 1.0 models in this study.

| parameter definition | parameter name | parameter value | source / reference |
|---|---|---|---|
| time frame of the simulation (years) | years.sim | 25 | 25 years: 1980-2005 |
| duration of acute stage | hiv.strain1.acute.dur.min<br>hiv.strain1.acute.dur.max | 12 | 12 weeks [19] |
| minimum time to AIDS | hiv.strain1.aids.progr.start.time | 104 | AIDS might start after 2 years (104 weeks) (only ~1% of infected people develop AIDS in the first 2 years) [20,22] |
| mean time to AIDS | hiv.strain1.aids.progr.mean.time | 468 | ~9 years (468 weeks) [20,22] |
| time to death when having AIDS | hiv.strain1.aids.dur | 52 | ~1 year (52 weeks) [20,22] |
| HSV-2 transmission – influence of gender | hsv2.tr.prob.f.m.ratio | 0.5 | [23,24] |
| HIV seed in 1980 | hiv.seed.strain1.initial.general | 0.005<br>for fitting: constant seed<br>100 simulations: seed drawn from a binomial distribution with probability = 0.005 | leads to an HIV prevalence of 0.6% for males and 1.2% for females in 1989 (values in Table S3) |
| HSV2 seed in 1980 | hsv2.seed.initial.general | 0.25 | [25] |

##### 4.11. Converting transmission probability parameters to hazard parameters

While in the majority of the literature, and also in StepSyn 1.0, transmission parameters are described as probabilities, the parameters of Simpact Cyan 1.0 are described in terms of hazards. Transmission probability parameters were converted to hazard parameters using the following formula [26]:

$$F(t)=1-\exp\left(-\int_0^t \lambda(x)dx\right)$$

where F(t) is the cumulative distribution function and λ(x) is the hazard function.

An example of conversion of a transmission probability parameter to a hazard parameter is presented below.

###### *4.11.1. Example*

Simpact Cyan 1.0 parameters:
put a = person.hsv2.a.dist.fixed.value and c = hsv2transmission.hazard.c
StepSyn parameters:
hsv2.tr.prob.chronic.m2f corresponds with a + c in Simpact Cyan 1.0
The product hsv2.tr.prob.chronic.m2f * hsv2.tr.prob.f.m.ratio corresponds with a in Simpact Cyan 1.0

<u>Calculation of a</u>
t=1/52 (because Simpact Cyan 1.0 is in years and StepSyn in weeks)
λ(x)=exp(a) (see Simpact Cyan 1.0 documentation)
F(t)=F(1/52)= hsv2.tr.prob.chronic.m2f * hsv2.tr.prob.f.m.ratio = 0.003*0.5 = 0.0015
If we fill in the equation

$$F(t)=1-\exp\left[-\int_0^t \lambda(x)dx\right]$$

We get

$$0.0015 = 1 - \exp\left[-\int_0^{1/52} \exp(a)dx\right]$$

$$-0.9985 = -\exp\left[-\int_0^{1/52} \exp(a)dx\right]$$

$$0.9985 = \exp\left[-\exp(a)\int_0^{1/52} dx\right]$$

$$\ln(0.9985) = -\exp(a)\cdot\frac{1}{52}$$

$$-52\cdot\ln(0.9985) = \exp(a)$$

$$a = \ln(-52\cdot\ln(0.9985)) = -2.550$$

So person.hsv2.a.dist.fixed.value = -2.550

Calculation of a + c

t=1/52 (because Simpact Cyan 1.0 is in years and StepSyn in weeks)

λ(x)=exp(a+c) (see Simpact Cyan 1.0 documentation)

F(t)=F(1/52)= hsv2.tr.prob.chronic.m2f = 0.003

$$0.003 = 1 - \exp\left[-\int_{0}^{1/52} \exp\,(a+c)dx\right]$$

In the same way as for a, we get

$a + c = \ln(-52 \cdot \ln(0.997)) = -1.856$

$c = -1.856 - a = -1.856 + 2.550 = 0.694$

So hsv2transmission.hazard.c = 0.694

#### 5. Results of model calibration

##### 5.1. Fit to HIV prevalence data (calibration targets)

Figure S12 compares the median HIV prevalence and the range of 100 simulations with the fitted parameters to the estimated HIV-1 prevalence data for Yaoundé's men and women in Table S3.

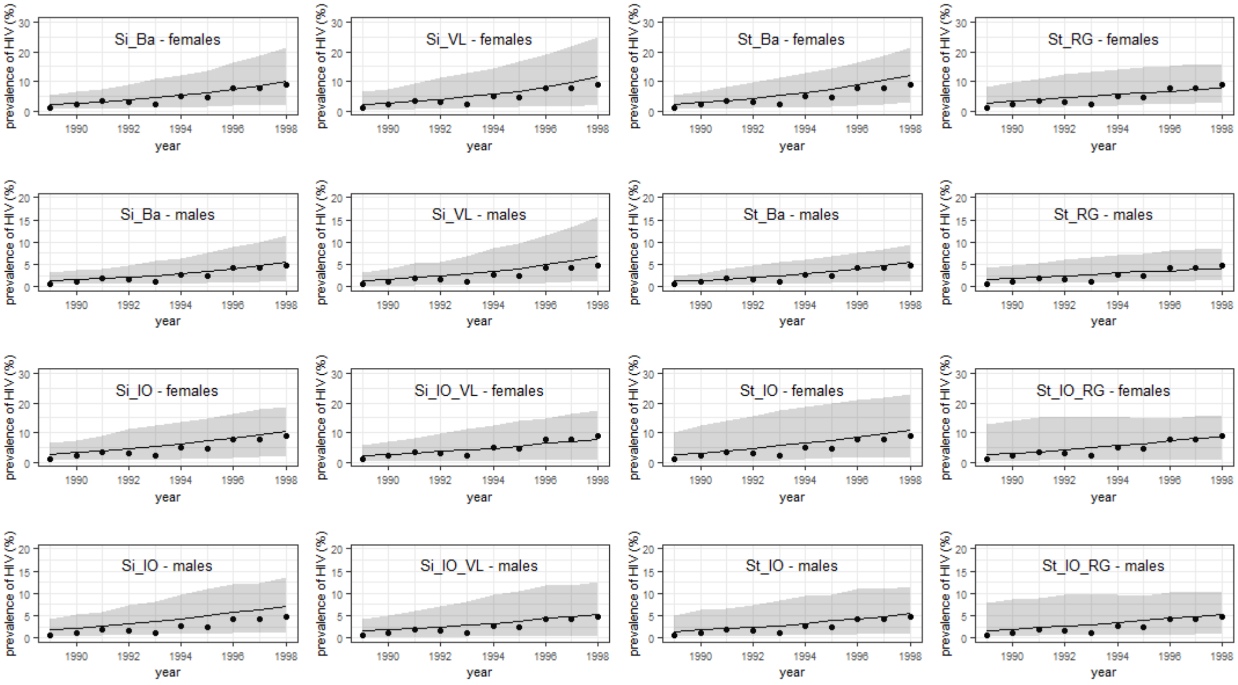


**Figure S12.** Median HIV prevalence (in %) of 100 simulations with the fitted parameters (solid line) and range ([minimum,maximum])(shaded area) for the period 1989–1998 (9–18 years into the simulation). The points represent the estimated HIV-1 prevalence data for Yaoundé's men and women in Table S3. Models: Si_Ba: Simpact 1.0 basic model; Si_IO: Simpact 1.0 model with inflow and outflow; Si_VL: Simpact 1.0 model with VL-dependent HIV transmission hazard; Si_IO_VL: Simpact 1.0 model with inflow, outflow, VL-dependent HIV transmission hazard; St_Ba: StepSyn 1.0 basic model; St_IO: Stepsyn 1.0 model with inflow and outflow; St_RG: StepSyn 1.0 model with STI life history explicitly modelled; St_IO_RG: StepSyn 1.0 model with inflow, outflow and STI life history explicitly modelled.

#### 5.2. HSV-2 prevalence curves

Figure S13 shows the median HSV-2 for females and the range of 100 simulations for the period 1980-2005 for the 8 models.

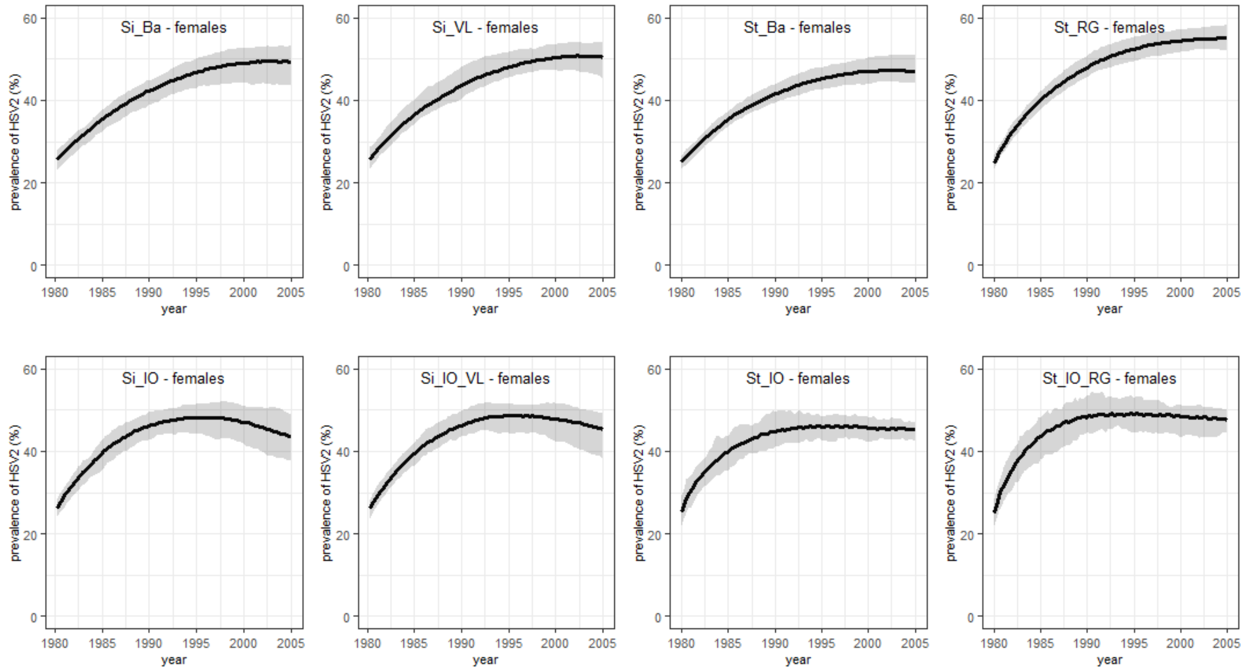


**Figure S13.** Median HSV2 prevalence for females (in %) of 100 simulations with the fitted parameters (solid line) and range ([minimum,maximum])(shaded area) for the period 1980–2005. Models: Si_Ba: Simpact 1.0 basic model; Si_IO: Simpact 1.0 model with inflow and outflow; Si_VL: Simpact 1.0 model with VL-dependent HIV transmission hazard; Si_IO_VL: Simpact 1.0 model with inflow, outflow, VL-dependent HIV transmission hazard; St_Ba: StepSyn 1.0 basic model; St_IO: Stepsyn 1.0 model with inflow and outflow; St_RG: StepSyn 1.0 model with STI life history explicitly modelled; St_IO_RG: StepSyn 1.0 model with inflow, outflow and STI life history explicitly modelled.

#### 6. Behavioural interventions

For the Simpact Cyan 1.0 models, the parameters describing the weight for the number of relationships a person already has ($\alpha_{numrel,man}$ and $\alpha_{numrel,woman}$ in formula (1) (parameter name in Simpact Cyan 1.0: formation.hazard.agegap.numrel_man, formation.hazard.agegap.numrel_woman) was changed. For the StepSyn 1.0 models, we changed the parameter for the proportion of male pending short links (concurrent unstable relationships) that are fulfilled to reach the individual preferred degree (parameter name in StepSyn 1.0: pending.short.links.fulfilled)(see Supplementary Material, section 1.2.2. for more detail).

Because sexual networks for Simpact Cyan 1.0 and StepSyn 1.0 are generated in different ways, the initial sexual network before applying interventions was different between Simpact Cyan 1.0 and StepSyn 1.0 models.

Figure S13 shows the distribution of the number of partners at the start of the HIV epidemic for both Simpact Cyan 1.0 and StepSyn 1.0 in case no intervention was implemented. In Simpact Cyan 1.0 the parameter for the weight for the number of relationships a person already has is equal to 0. In StepSyn 1.0 the parameter for the proportion of male pending short links that are fulfilled is equal to 1.

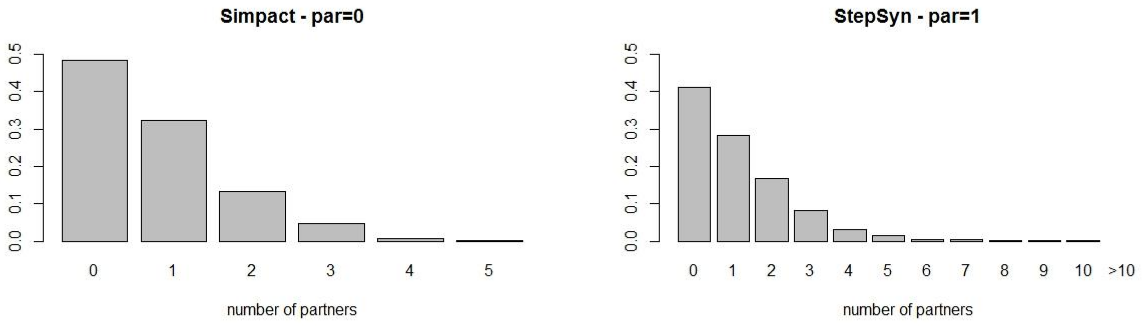


**Figure S14.** Distribution of the number of partners for the whole population at the start of the HIV epidemic (HIV seeding) for a simulation with Simpact Cyan 1.0 (left) and a simulation with StepSyn 1.0 (right).

We explored different values for the behavioural parameters for Simpact Cyan 1.0 and StepSyn 1.0 to determine a behavioural intervention that shows similar relative changes in the distribution of the number of partners. Tables S23 and S24 give a few examples for Simpact Cyan 1.0 and StepSyn 1.0 respectively.

**Table S23.** Mean, median, 75% percentile and 95% percentile of the distribution of the number of partners for different values of the parameter for the weight for the number of relationships a person already has in Simpact Cyan 1.0.

| parameter value | mean | median | 75% percentile | 95% percentile |
|---|---|---|---|---|
| 0 | 0.78 | 1 | 1 | 3 |
| -0.03 | 0.78 | 1 | 1 | 2 |
| -0.05 | 0.73 | 1 | 1 | 2 |
| -0.09 | 0.72 | 1 | 1 | 2 |

**Table S24.** Mean, median, 75% percentile and 95% percentile of the distribution of the number of partners for different values of the parameter for the proportion of male pending short links that are fulfilled in StepSyn 1.0.

| parameter value | mean | median | 75% percentile | 95% percentile |
|---|---|---|---|---|
| 1 | 1.16 | 1 | 2 | 4 |
| 0.7 | 1.09 | 1 | 2 | 3 |
| 0.4 | 0.98 | 1 | 1 | 3 |
| 0.1 | 0.78 | 1 | 1 | 2 |

Changing the weight for the number of relationships a person already has from 0 to -0.05 in Simpact Cyan 1.0, and changing the proportion of male pending short links that are fulfilled from 1 to 0.7 in StepSyn 1.0 reduces the mean number of partners by 6% and the 95% percentile with 1, while keeping the median and the 75% percentile unchanged.

For all models, behavioural interventions also reduced the median HSV-2 prevalence (see Table S25).

**Table S25.** Results for the predicted HSV-2 prevalence in 1997 for females and males in case of no behavioural intervention, a behavioural intervention implemented in 1990, and lower promiscuity from 1980 onwards. Median HIV prevalence (in %) and range ([minimum,maximum]) of 100 simulations.

| | No intervention | | Intervention in 1990 | | Lower promiscuity from 1980 onwards | |
|---|---|---|---|---|---|---|
| Model | Females | Males | Females | Males | Females | Males |
| Si_Ba | 48.0 (43.6-51.5) | 38.2 (34.8-41.6) | 47.4 (44.0-51.8) | 37.9 (35.6-42.1) | 45.8 (41.7-49.5) | 36.9 (34.5-39.6) |
| Si_IO | 48.0 (43.1-51.6) | 38.6 (34.7-42.4) | 47.7 (43.8-51.5) | 38.2 (35.1-40.7) | 47.3 (42.4-49.9) | 37.9 (34.3-41.1) |
| Si_VL | 49.3 (46.7-52.4) | 38.8 (36.1-42.9) | 48.7 (45.4-51.9) | 38.6 (35.9-42.4) | 47.4 (44.0-50.1) | 37.6 (34.7-41.3) |
| Si_IO_VL | 48.7 (44.3-51.2) | 38.8 (34.3-43.0) | 47.9 (44.0-51.5) | 38.6 (35.4-42.0) | 47.3 (43.6-50.9) | 37.7 (35.1-41.1) |
| St_Ba | 46.3 (43.5-49.0) | 37.0 (35.2-38.9) | 45.6 (43.0-48.1) | 36.7 (34.8-38.8) | 43.1 (40.2-45.3) | 35.4 (33.3-36.7) |
| St_IO | 45.9 (43.1-48.7) | 35.3 (32.9-37.6) | 43.4 (40.8-46.5) | 34.1 (31.7-35.7) | 42.3 (39.1-45.5) | 33.5 (31.2-35.5) |
| St_RG | 53.7 (50.6-56.3) | 40.4 (38.2-41.7) | 52.4 (50.3-55.4) | 39.7 (37.6-41.6) | 48.4 (45.8-51.5) | 37.5 (36.1-39.5) |
| St_IO_RG | 48.7 (44.5-52.4) | 36.4 (32.1-38.8) | 45.1 (42.3-48.7) | 34.4 (32.9-36.2) | 44.4 (41.7-46.9) | 34.0 (32.0-36.2) |

Models: Si_Ba: Simpact 1.0 basic model; Si_IO: Simpact 1.0 model with inflow and outflow; Si_VL: Simpact 1.0 model with VL-dependent HIV transmission hazard; Si_IO_VL: Simpact 1.0 model with inflow, outflow, VL-dependent HIV transmission hazard; St_Ba: StepSyn 1.0 basic model; St_IO: Stepsyn 1.0 model with inflow and outflow; St_RG: StepSyn 1.0 model with STI life history explicitly modelled; St_IO_RG: StepSyn 1.0 model with inflow, outflow and STI life history explicitly modelled.

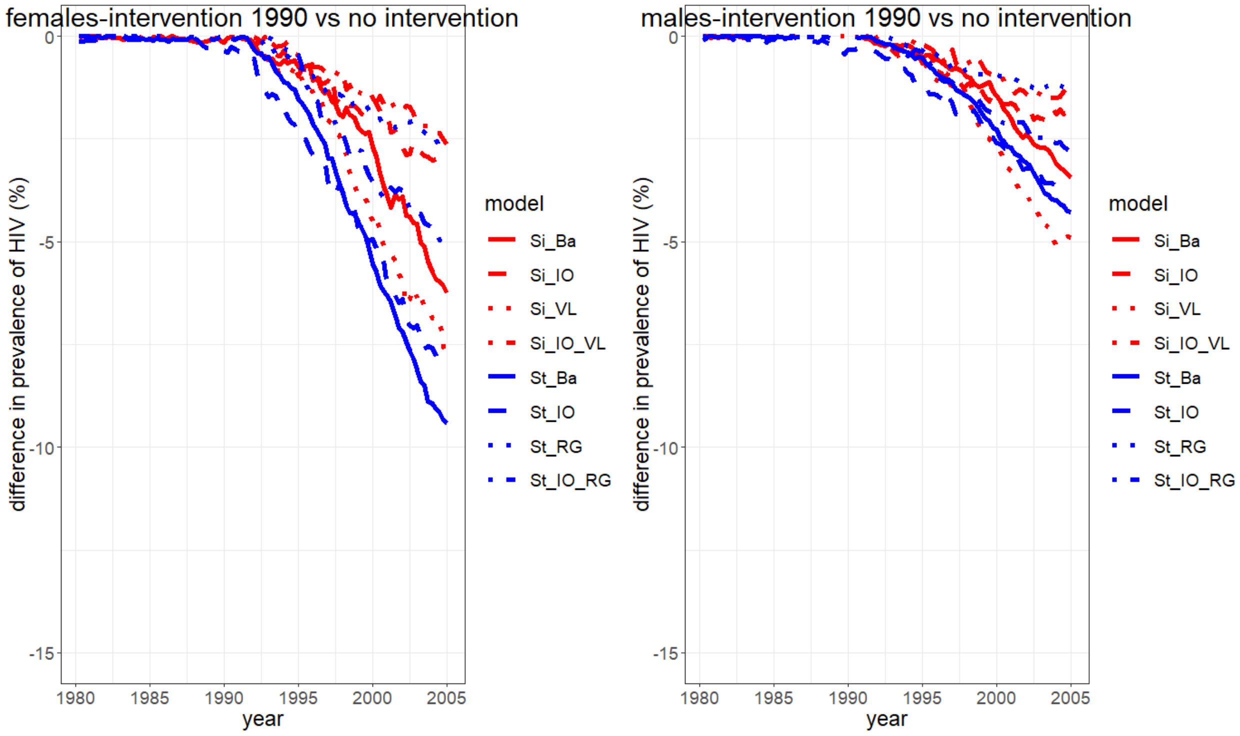


**Figure S15.** Decrease in HIV prevalence in case of an intervention in 1990, compared to the scenario without intervention. Median HIV prevalence (in %) of 100 simulations. Left: females; right: males. Models: Si_Ba: Simpact 1.0 basic model; Si_IO: Simpact 1.0 model with inflow and outflow; Si_VL: Simpact 1.0 model with VL-dependent HIV transmission hazard; Si_IO_VL: Simpact 1.0 model with inflow, outflow and VL-dependent HIV transmission hazard; St_Ba: StepSyn 1.0 basic model; St_IO: Stepsyn 1.0 model with inflow and outflow; St_RG: StepSyn 1.0 model with STI life history explicitly modelled; St_IO_RG: StepSyn 1.0 model with inflow, outflow and STI life history explicitly modelled.

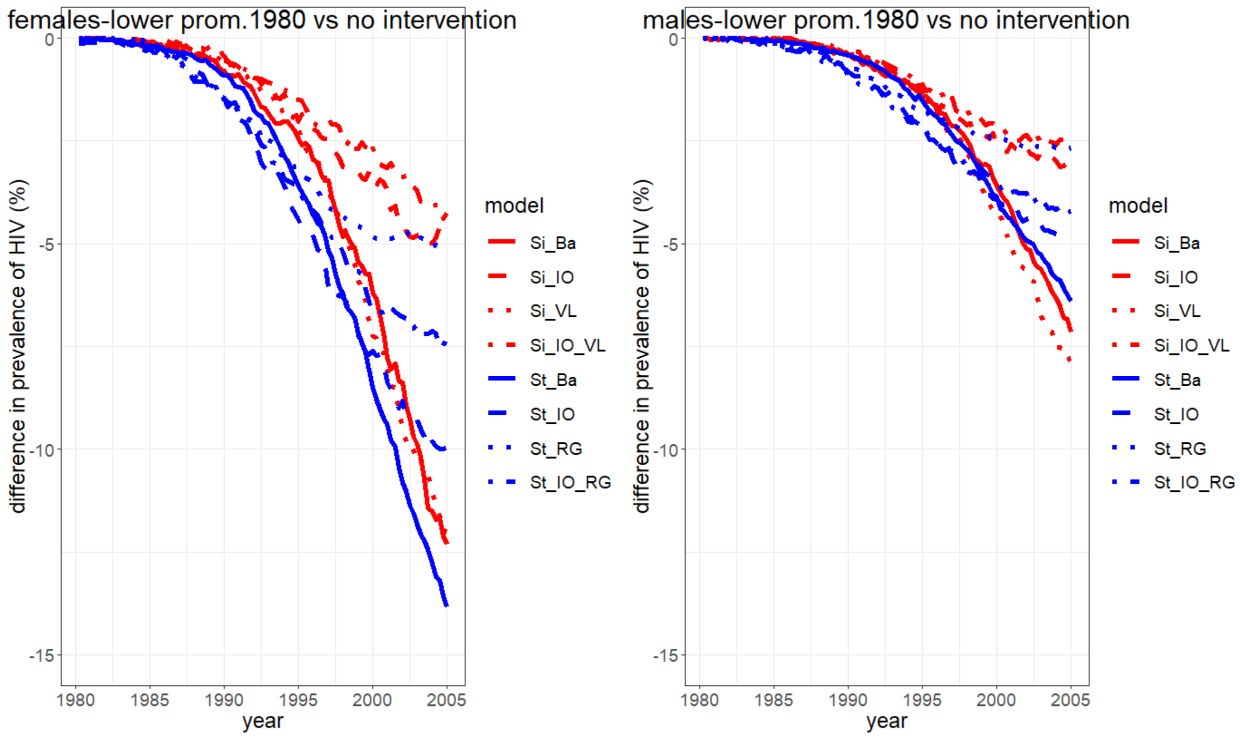


**Figure S16.** Decrease in HIV prevalence in case of lower promiscuity in 1980, compared to the scenario without intervention. Median HIV prevalence (in %) of 100 simulations. Left: females; right: males. Models: Si_Ba: Simpact 1.0 basic model; Si_IO: Simpact 1.0 model with inflow and outflow; Si_VL: Simpact 1.0 model with VL-dependent HIV transmission hazard; Si_IO_VL: Simpact 1.0 model with inflow, outflow and VL-dependent HIV transmission hazard; St_Ba: StepSyn 1.0 basic model; St_IO: Stepsyn 1.0 model with inflow and outflow; St_RG: StepSyn 1.0 model with STI life history explicitly modelled; St_IO_RG: StepSyn 1.0 model with inflow, outflow and STI life history explicitly modelled.
